# Endothelial SIRT3 regulates myofibroblast metabolic shifts in diabetic kidneys

**DOI:** 10.1101/2020.12.15.422824

**Authors:** Swayam Prakash Srivastava, Jinpeng Li, Yuta Takagaki, Munehiro Kitada, Julie Goodwin, Keizo Kanasaki, Daisuke Koya

## Abstract

Defects in endothelial cells cause deterioration in kidney function and structure. Here, we found that endothelial SIRT3 regulates metabolic reprogramming and fibrogenesis in the kidneys of diabetic mice. By analyzing, gain-of-function of the SIRT3 gene by overexpression in a fibrotic mouse strain conferred disease resistance against diabetic kidney fibrosis; while its loss-of-function in endothelial cells exacerbated the levels of diabetic kidney fibrosis. Regulation of endothelial cell SIRT3 on fibrogenic processes was due to tight control over the defective central metabolism and linked-activation of endothelial-to-mesenchymal transition (EndMT). SIRT3 deficiency in endothelial cells stimulated the TGFβ/Smad3-depandent mesenchymal transformations in renal tubular epithelial cells. These data demonstrate that SIRT3 regulates defective metabolism and EndMT-mediated activation of the fibrogenic pathways in the diabetic kidneys. Together, our findings show that endothelial cell SIRT3 is a fundamental regulator of defective metabolism regulating health and disease processes in the kidney.

**Context and significance:** The present study describes that endothelial cell SIRT3 protects against renal fibrosis by mitigating the metabolic reprogramming and associated mesenchymal transformations in diabetes. Accordingly, amelioration of endothelial cell SIRT3 and targeting the defective metabolism offer a potential therapeutic approach to treat or prevent the progression of diabetic kidney disease. Further investigation would delineate the mechanisms of SIRT3 up-regulation in diabetic kidney disease and discover small molecules to increase the expression of the SIRT3. In addition, this study describes the biology of SIRT3 in the health and disease processes of kidney endothelial cells.

## Introduction

Of over 400 million people with diabetes, about one-third will develop diabetic kidney disease (DKD), a leading cause of end-stage renal disease (ESRD) that causes more than 950,000 deaths each year globally (Badal and Danesh, 2014; Cooper and Warren, 2019; Reidy et al., 2014). Over the last two decades, no new drugs have been approved to specifically prevent DKD or to improve kidney functions (Breyer and Susztak, 2016). This lack of progress likely stems from poor understanding of the mechanism of kidney dysfunction and this knowledge gap contributes to the suboptimal treatment options available for these patients. Improved understanding of the mechanism of pathogenesis of diabetic kidney disease is urgently needed to catalyze the development of novel therapeutics which can be targeted to the early stages of these diseases, before kidney damage becomes irreversible.

Although kidney fibrosis is identical manifestation in all progressive form of chronic kidney disease and DKD, and is caused by excess deposition of extracellular matrix leading to renal function deterioration, and kidney injury (LeBleu et al., 2013; Srivastava et al., 2019; Zeisberg et al., 2003). Fibroblasts play a crucial role in kidney fibrosis but the origin of fibroblasts is still obscure (Grande and Lopez-Novoa, 2009; LeBleu et al., 2013; Srivastava et al., 2013; Zeisberg and Neilson, 2010). There are six well-reported sources of matrix-producing myofibroblasts: (1) activated residential fibroblasts, (2) differentiated pericytes, (3) recruited circulating fibrocytes, (4) from mesenchymal cells derived from M2 phenotype macrophages via macrophage-to-mesenchymal transition from mesenchymal cells derived from tubular epithelial cells via epithelial-to-mesenchymal transition (EMT) and (6) from mesenchymal cells transformed from endothelial cells (ECs) via endothelial-to-mesenchymal transition (EndMT) (Grande and Lopez-Novoa, 2009; Grande et al., 2015; LeBleu et al., 2013; Srivastava et al., 2019; Wang et al., 2017). Among these diverse origins of matrix-producing fibroblasts, EndMT is an important source of myofibroblasts in several organs, including in kidney (Medici, 2016; Srivastava et al., 2019). EndMT is characterized by the loss of endothelial markers, including cluster of differentiation 31 (CD31), and acquisition of the expression of mesenchymal proteins including a-smooth muscle actin (αSMA), fibronectin and SM22α (Srivastava et al., 2019). The complete conversion from endothelial cells into mesenchymal cell types is not essential; although intermediate phenotypes are sufficient to induce alteration in fibrogenic programs (Kizu et al., 2009; LeBleu et al., 2013; Zeisberg et al., 2007).

Endothelial cells are key players in the formation of new blood vessels both in health and life-threatening diseases (Eelen et al., 2015; Eelen et al., 2018). 6-phosphofructo-2-kinase/fructose-2,6-bisphosphatase-3 (PFKFB3)-driven glycolysis regulate the endothelial cell metabolism and vessel sprouting, whereas carnitine palmitoyltransferase 1a (CPT1a)-mediated fatty acid oxidation regulates the proliferation of endothelial cells in the stalk of the sprout (Cantelmo et al., 2016; Cruys et al., 2016; De Bock et al., 2013; Schoors et al., 2015; Schoors et al., 2014). To maintain vascular homeostasis, endothelial cells use metabolites for epigenetic regulation of endothelial cell sub-type differentiation and maintain crosstalk through metabolite release with other cell types (Eelen et al., 2018; Schoors et al., 2015). Disruption of metabolic homeostasis in endothelial cells contributes to disease phenotypes (Theodorou and Boon, 2018; Zhou et al., 2019). Importantly, EndMT causes alteration of endothelial cell structure and metabolism, which is an area of active investigation (Chen et al., 2012; Lovisa and Kalluri, 2018; Xiong et al., 2018). The mesenchymal cells derived from EndMT reprogram their metabolism and depend on glycolytic metabolites for nucleic acid, amino acids, glycoproteins, and lipid synthesis (Eelen et al., 2018; Lovisa and Kalluri, 2018; Theodorou and Boon, 2018; Xiong et al., 2018). Researchers have examined the contribution of EndMT to renal fibrosis in several mouse models of chronic kidney disease (Grande and Lopez-Novoa, 2009; LeBleu et al., 2013; Liu, 2011; Medici and Kalluri, 2012; Srivastava et al., 2014; Zeisberg et al., 2008; Zeisberg and Neilson, 2010). Zeisberg et al., performed seminal experiment and reported that approximately 30~50% of fibroblasts co-expressed the endothelial marker CD31 and markers of fibroblasts and myofibroblasts such as fibroblast specific protein-1 (FSP-1) and αSMA in the kidneys of mice experienced to unilateral ureteral obstructive nephropathy (UUO) (Zeisberg et al., 2008). EndMT contributes to the accumulation of activated fibroblasts; thus, targeting EndMT might have therapeutic potential against renal fibrosis (LeBleu et al., 2013; Li et al., 2020; Liu, 2011; Medici and Kalluri, 2012; Srivastava et al., 2020a; Zeisberg et al., 2008).

A correlation between mitochondrial damage, inflammation and, renal fibrosis has been demonstrated suggesting, that mitochondrial integrity and mitochondrial metabolism is critical for kidney cells homeostasis (Chung et al., 2019). Mitochondrial sirtuins play a key role in the regulation of mitochondrial integrity and metabolism and during recent years the involvement of mitochondrial sirtuins are now been gaining momentum in kidney research (Hershberger et al., 2017; Morigi et al., 2015; Perico et al., 2016). Among mitochondrial sirtuins, SIRT3 is a major deacetylase that targets several diverse enzymes involved in central metabolism, resulting in the activation of many oxidative pathways (Yin and Cadenas, 2015). SIRT3 blocks the characteristics of organ fibrosis by regulating TGF-β/smad signaling (Chen et al., 2015; Sosulski et al., 2017; Sundaresan et al., 2015). SIRT3 regulates the abnormal glucose metabolism via tight control over PKM2 tetramer-to-dimer interconversion and HIF1α accumulation in the diabetic kidneys (Srivastava et al., 2018). SIRT3 deplete tubular epithelial cells are highly dependent on reprogrammed defective metabolism and are associated with higher mesenchymal activation in the diabetic kidneys (Srivastava et al., 2020b; Srivastava et al., 2018).

In the present study, we aimed to understand the contribution of endothelial cell SIRT3 in the regulation of metabolic reprograming and fibrogenic processes in the kidneys. Therefore, the development of suitable animal models for studying the functional and physiological implication of endothelial cell SIRT3 is critical to understand the pathogenesis of diabetic kidney disease. To begin to understand how endothelial cell SIRT3 may be regulating renal fibrosis in diabetes, we developed two unique novel mouse models 1) overexpression mouse model in fibrotic background, 2) endothelial specific deletion of SIRT3 gene in less-fibrotic mouse background.

Our results indicate a key role of endothelial cell SIRT3 in the regulation the metabolic reprogramming and-linked activation of EndMT processes, which contributes in fibrogenic phenotype in the kidneys of diabetic mice.

## Results

### SIRT3 deficiency in endothelial cells is a critical fibrogenic phenotype in the kidneys of diabetic mice

The streptozotocin (STZ)-induced diabetic CD-1 is the established mouse model to study diabetic kidney disease (Kanasaki et al., 2014; Shi et al., 2015; Srivastava et al., 2016). In mice, the kidney fibrosis phenotype is largely dependent upon the strain specificity (Srivastava et al., 2016). The STZ-induced diabetic CD-1 mice and diabetic C57BL6 mouse experienced similar level of blood glucose however, the kidneys of diabetic CD-1 mice have been shown to display higher rate of EndMT and massive fibrosis when compared to the kidneys of diabetic C57Bl6 mice (Kanasaki et al., 2014; Srivastava et al., 2018). Here, we confirmed the dose dependent effect of STZ in the development of renal fibrosis in the CD-1 and C57Bl6 mouse strains however, the kidneys of diabetic CD-1 mice experienced higher fibrosis when compared to diabetic C57Bl6 mice **(Figure S1a-b)**. Therefore, diabetic CD-1 mouse is known as fibrotic strain however, the diabetic C57Bl6 mouse is considered as less-fibrotic strain (Srivastava et al., 2018; Sugimoto et al., 2007). The kidneys of diabetic CD-1 mice displayed complete suppression in SIRT3 protein when compared to non-diabetic control, whereas, diabetic C57Bl6 did not **(Figure 1a)**. Multiplex immunohistochemical analysis of vimentin/aminopeptidase A (a marker of proximal tubule) or vimentin/uromodulin (a marker of distal tubule) showed higher level of vimentin in proximal and distal tubules in the kidneys of diabetic CD-1 mice when compared to non-diabetic control and such effects were not prominent in kidneys of diabetic C57Bl6 mice (**Figure 1b)**. Furthermore, we found that CD31-positive cells in diabetic CD-1 mouse kidneys displayed significant suppression of SIRT3 whereas the kidneys of diabetic C57Bl6 mice did not reveal a remarkable difference in SIRT3 protein levels **(Figure 1c)**. In addition, we found that the endothelial cells isolated from the kidneys of diabetic CD-1 mice, showed significant suppression in SIRT3 and CD31 protein level **(Figure 1d)**.

**Figure 1.**
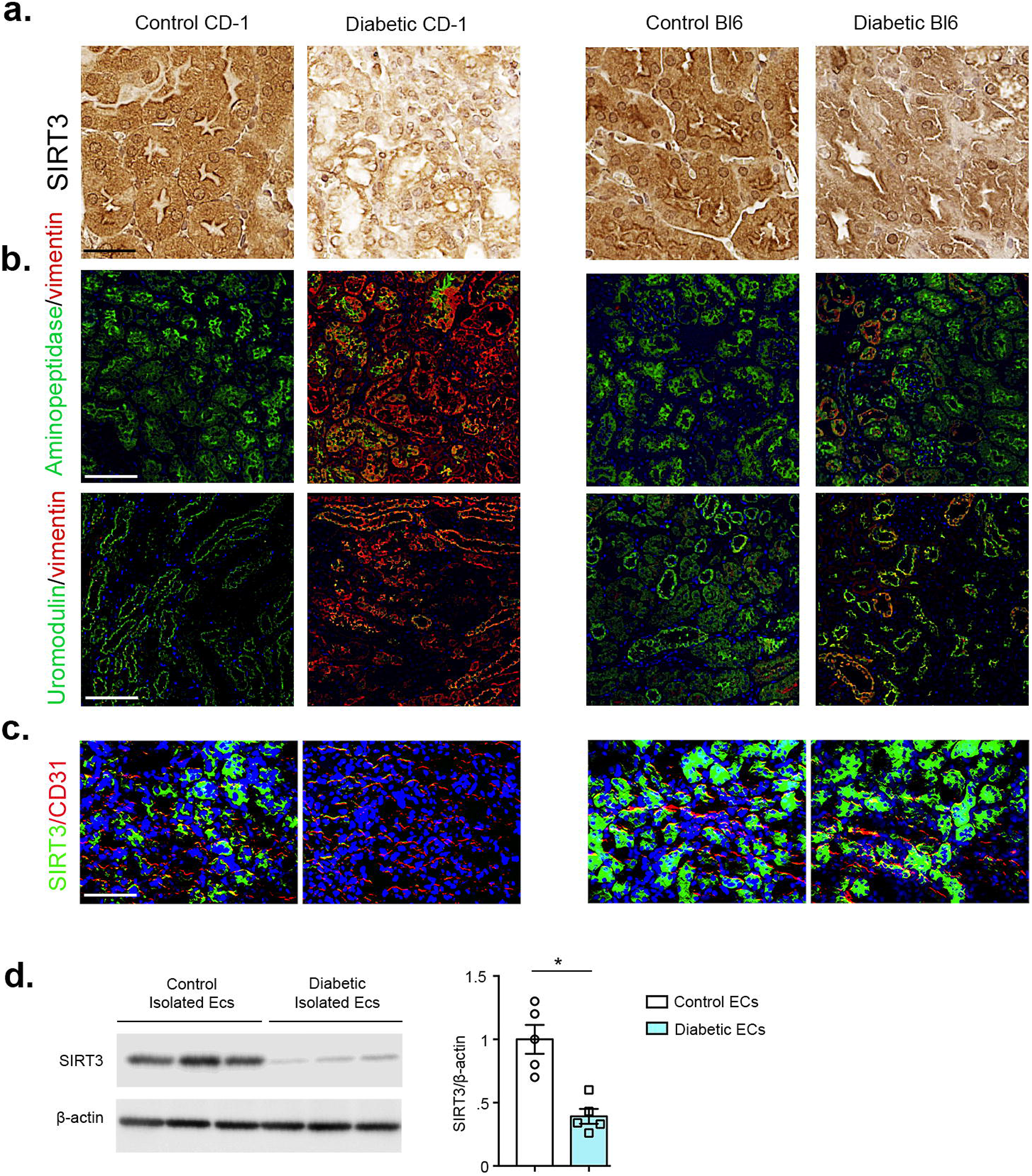
Diabetic kidney disease is associated with suppression of endothelial SIRT3 protein. **(a)** Immunohistochemical analysis of the kidneys of control and diabetic CD-1, and C57Bl6 mice. Representative pictures are shown. 40X images are shown. N=7/each group. Scale bar: 50 μm. **(b)** Immunofluorescent analysis for aminopeptidase A/vimentin and uromodulin/Vimentin in the kidneys of control and diabetic mice of CD-1 and C57Bl6 mice. Representative pictures are shown. N=7/each group. Scale bar: 50 μm. **(c)** Immunofluorescent analysis of the kidneys of control and diabetic mice of CD-1 and C57Bl6 mice. FITC-labeled SIRT3, Rhodamine-labeled CD31 and DAPI blue. Scale bar: 50 μm. Representative pictures are shown. 40X images are shown. N=7/CD1 mice whereas N=5/C57Bl6 mice were analyzed. Data in the graph are shown as mean ± SEM. **(d)** Western blot analysis of SIRT3 protein in isolated endothelial cells from the kidneys of control and diabetic CD-1 mice. Densitometry analysis was normalized by β-actin. N=5 were analyzed in each group. Representative blots are shown. Data in the graph are shown as mean ± SEM.

### SIRT3 over-expression in fibrogenic phenotype protects from diabetes-associated renal fibrosis

Our general approach was to define how endogenous SIRT3 in the endothelial cell was associated to repress the fibrogenic characteristics in diabetic kidney? To answer, we bred SIRT3 over-expression mouse in the fibrogenic mouse background (CD-1 mouse background). We back-crossed Tie1-Sirt3 tg mice (C57Bl6 background that displayed expressed levels of SIRT3 protein in the endothelial cells) with CD-1 mice. The purpose of back-cross breeding was to transfer the Sirt3 Tg gene (which is in less fibrotic C57Bl6 background) to the endothelial cells of CD-1 mouse (fibrotic phenotype). The schematic diagram depicts the back-cross scheme between CD-1 and Sirt3 tg mice **(Figure 2a)**. At the 9^th^ generation, SIRT3 mRNA/protein level was significantly upregulated in the isolated ECs from the kidneys of SIRT3 overexpressed mouse Sirt3 Tg (+); CD-1 (from here denoted eEx) when compared to Sirt3 Tg-; CD-1 (control) mice **(Fig. 2b-c)**. We injected a single higher dose of STZ (200 mg/kg/day i.p.) in the eEx and littermate control. At the time of sacrifice, non-diabetic eEX and non-diabetic controls had similar body weight, blood glucose, kidney weight, albumin-to-creatinine ratio (ACR), blood pressure, liver weight, and heart weight, However, diabetic eEx had relatively lower kidney weight and significantly suppressed ACR levels when compared to diabetic controls **(Figure S2)**. We did not observe any remarkable differences in the body weight, blood glucose, blood pressure, liver weight, or heart weight in the diabetic eEx when compared to diabetic controls **(Figure S2).** We observed minor fibrotic alterations between non-diabetic control and non-diabetic eEx; however, diabetic eEx exhibited significantly lower levels of relative area fibrosis (RAF), relative collagen deposition (RCD) and glomerular surface area when compared to diabetic controls **(Figure 2d and Figure S3)**. The diabetic kidneys of female mice displayed less fibrotic alterations when compared to diabetic kidneys of male mice; moreover, the kidneys of diabetic female eEX mice displayed suppressed level of fibrosis when compared to the kidneys of diabetic female control mice **(Figure 2d and Figure S4)**. Multiplex immunohistochemical data of aminopeptidase/⍰-SMA and uromodulin/⍰-SMA revealed a significant reduction in ⍰-SMA level in the proximal tubules and in distal tubules in the kidneys of diabetic eEx **(Figure 2e)**. The kidneys of diabetic eEx mice had suppressed protein level of collagen I, fibronectin, vimentin and ⍰-SMA when compared to the kidneys of diabetic control **(Figure 2f)**.

**Figure 2.**
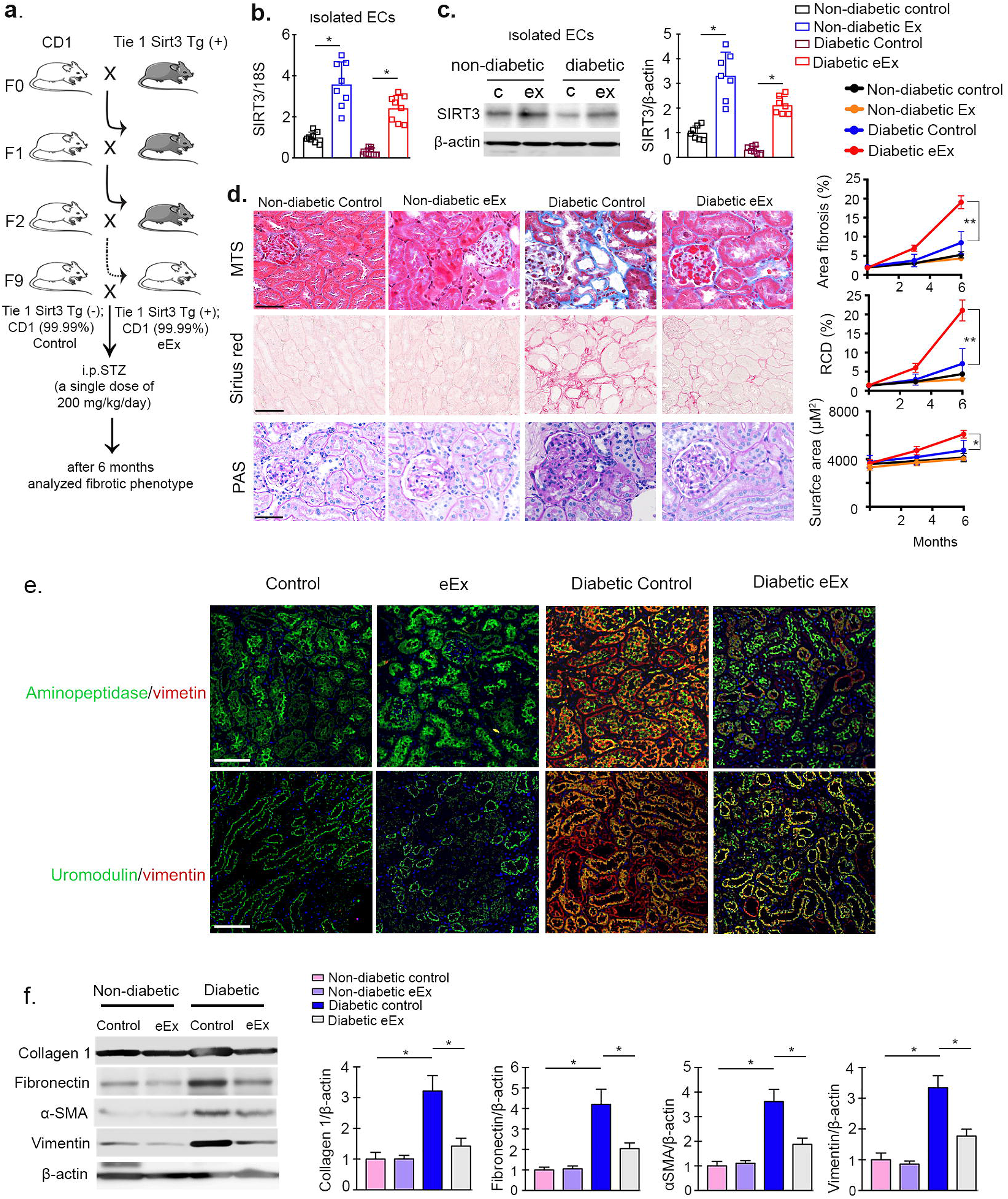
Over-expression of endothelial SIRT3 protects against fibrosis in the kidney of diabetic mice. **(a)** Schematic chart of back crossing of endothelial specific Sirt3tg (Tie 1 Sirt3 tg+) with CD-1 mice. After the 9^th^ generation, 99.99% of the genetic background of CD-1 mice was transferred. A single dose of STZ (200 mg/kg/day i.p.) were injected in the control (Tie 1 Sirt3 tg-; CD-1) and eEx (Tie 1 Sirt3 tg +; CD-1) mice to induce fibrosis. **(b)** SIRT3 mRNA expression was analyzed by qPCR in the isolated endothelial cells of eEX and control littermates. 18S was used as internal control. N=8/each group. **(c)** Western blot analysis of SIRT3 protein in isolated endothelial cells from the kidney of control and eEx mice. N=7/each group. Representative blots are shown. Densitometry calculation were normalized to β-actin. **(d)** Masson trichrome, Sirius red and PAS staining in the kidney of nondiabetic and diabetic, eEx and control littermates. Representative images are shown. Area of fibrosis (%), relative collagen deposition (RCD %) and surface area (μm^2^) were measured using the ImageJ program. N=7/each group. Scale bar: 50 μm. 40X images in the MTS and PAS while 30x images in the Sirius red. Data in the graph are shown as mean ± SEM. **(e)** Immunofluorescent analysis for aminopeptidase A/αSMA and uromodulin/αSMA in the kidneys of nondiabetic and diabetic, control and eEx. Representative images are shown. Scale bar: 50 mm in each panel. N=7/each group. **(f)** Western blot analysis of collagen I, fibronectin, α-SMA and vimentin in the kidney of non-diabetic and diabetic, control and eEx mice. N=5/each group. Representative blots are shown. Densitometry calculation were normalized to β-actin. Data in the graph are shown as mean ± SEM.

### Loss of endothelial SIRT3 worsens diabetes-associated fibrosis in kidney

To study the loss of function of SIRT3, we deleted the *Sirt3* gene in endothelial cells. Crossing of VE cadherin–Cre mice with mice having floxed alleles of SIRT3, resulted in the excision of the *Sirt3* gene (SIRT3 fl/fl; VE cadherin Cre+; from here denoted eKO), leading to the absence of SIRT3 protein expression in endothelial cells of the kidney when compared to littermate controls (SIRT3 fl/fl; VE cadherin Cre-) which exhibit no recombinase activity. The schematic diagram depicts the generation of eKO mice **(Figure 3a)**. SIRT3 mRNA/protein levels were significantly diminished in the endothelial cells isolated from the kidneys of eKO mice **(Figure 3b-c)**. At the age of 10 weeks, we injected 5 consecutive multiple low doses of STZ (50 mg/kg/day i.p.) and after 4 months analyzed the level of fibrosis in the eKO and littermate control. At the time of sacrifice, non-diabetic eKO and non-diabetic littermate controls had no remarkable change in body weight, blood glucose, kidney weight, ACR, blood pressure, liver weight and heart weight; however, diabetic eKO had relatively higher kidney weight and significantly higher ACR when compared to diabetic littermate controls **(Figure S5).** We did not observe a remarkable difference in the body weight, blood glucose, blood pressure, liver weight, or heart weight in the diabetic eKO when compared to diabetic littermate controls **(Figure S5).** Minor fibrotic alterations between non-diabetic controls and non-diabetic eKO were observed; however, diabetic eKO exhibited higher relative area of fibrosis, relative collagen deposition and severe glomerulosclerosis when compared to diabetic littermate controls **(Figure 3d and Figure S6)**. The diabetic kidneys of female mice displayed less fibrotic alterations when compared to diabetic kidneys of male mice; moreover, the kidneys of diabetic female eKO mice displayed higher level of fibrosis when compared to the kidneys of diabetic female control mice **(Figure 3d and Figure S7).** Multiplex immunohistochemical data of aminopeptidase/⍰-SMA and uromodulin/⍰-SMA revealed a significant higher ⍰-SMA level in the proximal tubules and in distal tubules in the kidneys of diabetic eKO when compared to diabetic control **(Figure 3e)**. The kidneys of diabetic eKO mice had induced protein level of collagen I, fibronectin, vimentin and ⍰-SMA when compared to the kidneys of diabetic control **(Figure 3f)**.

**Figure 3.**
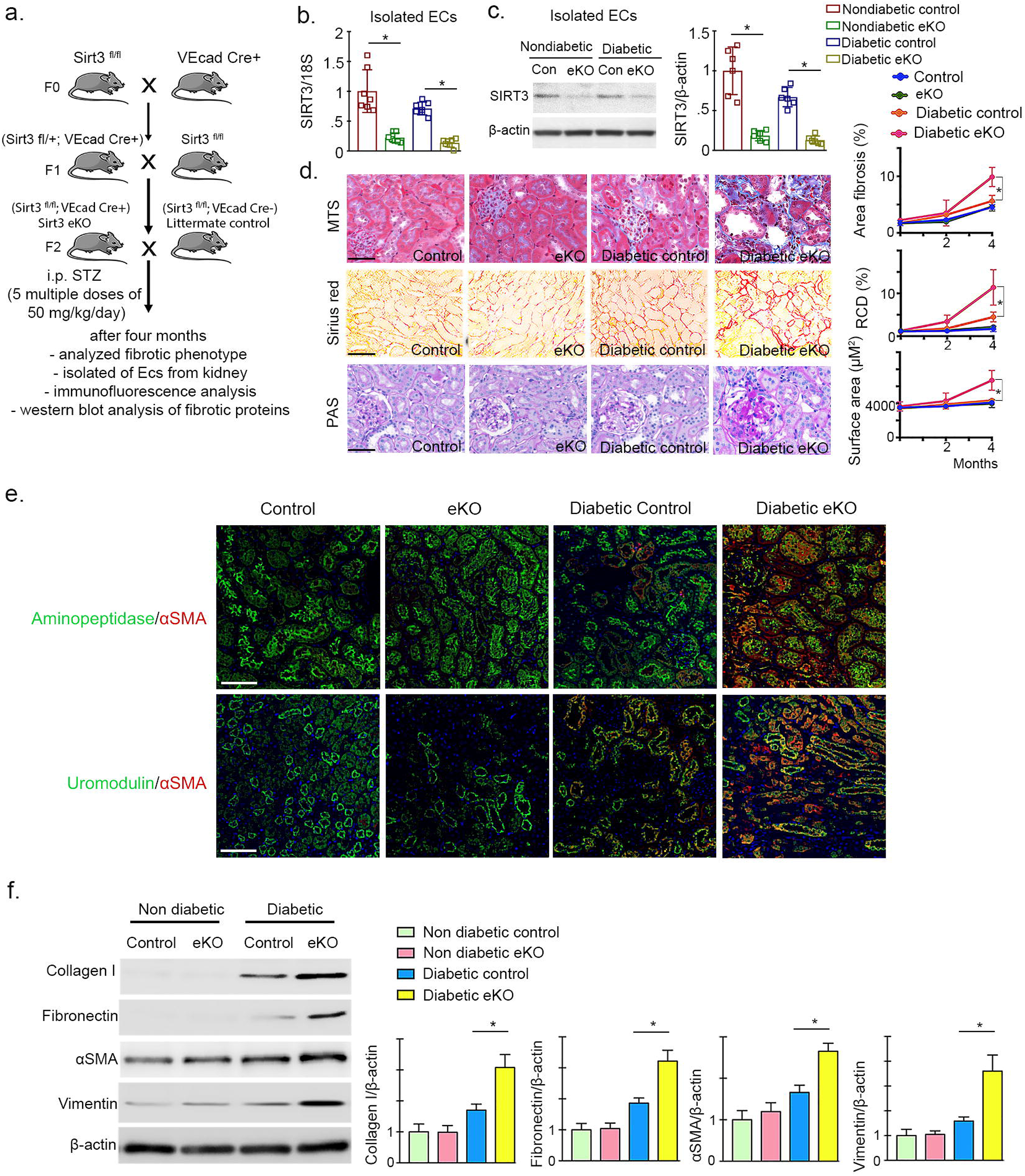
Loss of endothelial SIRT3 worsens renal fibrosis in the mouse model of diabetic kidney disease. **(a)** Schematic chart showing the generation of endothelial specific Sirt3 knockout mice. 5 multiple low doses of STZ (50 mg/kg/day i.p.) were injected in the control (Sirt3 ^fl/fl^; VeCad Cre-) and eKO (Sirt3 ^fl/fl^; VeCad Cre+) mice to induce fibrosis. **(b)** SIRT3 mRNA expression level was analyzed by qPCR in the isolated endothelial cells of eKO and control littermates. 18S was used as internal control. N=7/each group. **(c)** Western blot analysis of SIRT3 protein in isolated endothelial cells from the kidneys of control and eKO mice. N=6/each group. Representative blots are shown. **(d)** Masson trichrome, Sirius red and PAS staining in the kidneys of non-diabetic and diabetic, control littermates and eKO mice. Representative images are shown. Area of fibrosis (%), relative collagen deposition (RCD %) and surface area were measured using the ImageJ program. N=7/each group. Data in the graph are shown as mean ± SEM. Scale bar: 50 μm in MTS and PAS panel while 70 μm in Sirius red. 40X images in the MTS and PAS while 30x images in the Sirius red. **(e)** Immunofluorescent analysis for aminopeptidase A/αSMA and uromodulin/αSMA in the kidneys of nondiabetic and diabetic, control littermates and eKO mice. Representative images are shown. Scale bar: 50 mm in each panel. N=7/each group. **(f)** Western blot analysis of collagen I, fibronectin, α-SMA and vimentin in the kidney of non-diabetic and diabetic, control littermates and eKO mice. N=6/each group. Representative blots are shown. Densitometry calculation were normalized to β-actin.

### SIRT3 regulates endothelial-to-mesenchymal transitions in the kidneys

The kidneys of diabetic eEX exhibited suppressed levels of FSP-1 and αSMA in CD31 positive cells when compared to diabetic controls (severe fibrosis in kidneys), whereas the kidneys of diabetic eKO displayed significantly higher levels of FSP-1, αSMA and TGFβR1 in CD31 positive cells when compared to diabetic littermate controls (less fibrosis in kidneys) **(Figure 4a and b)**. However, there were no remarkable alterations in the levels of FSP-1 and αSMA in the kidneys of non-diabetic mice **(Figure 4a and b)**.

**Figure 4.**
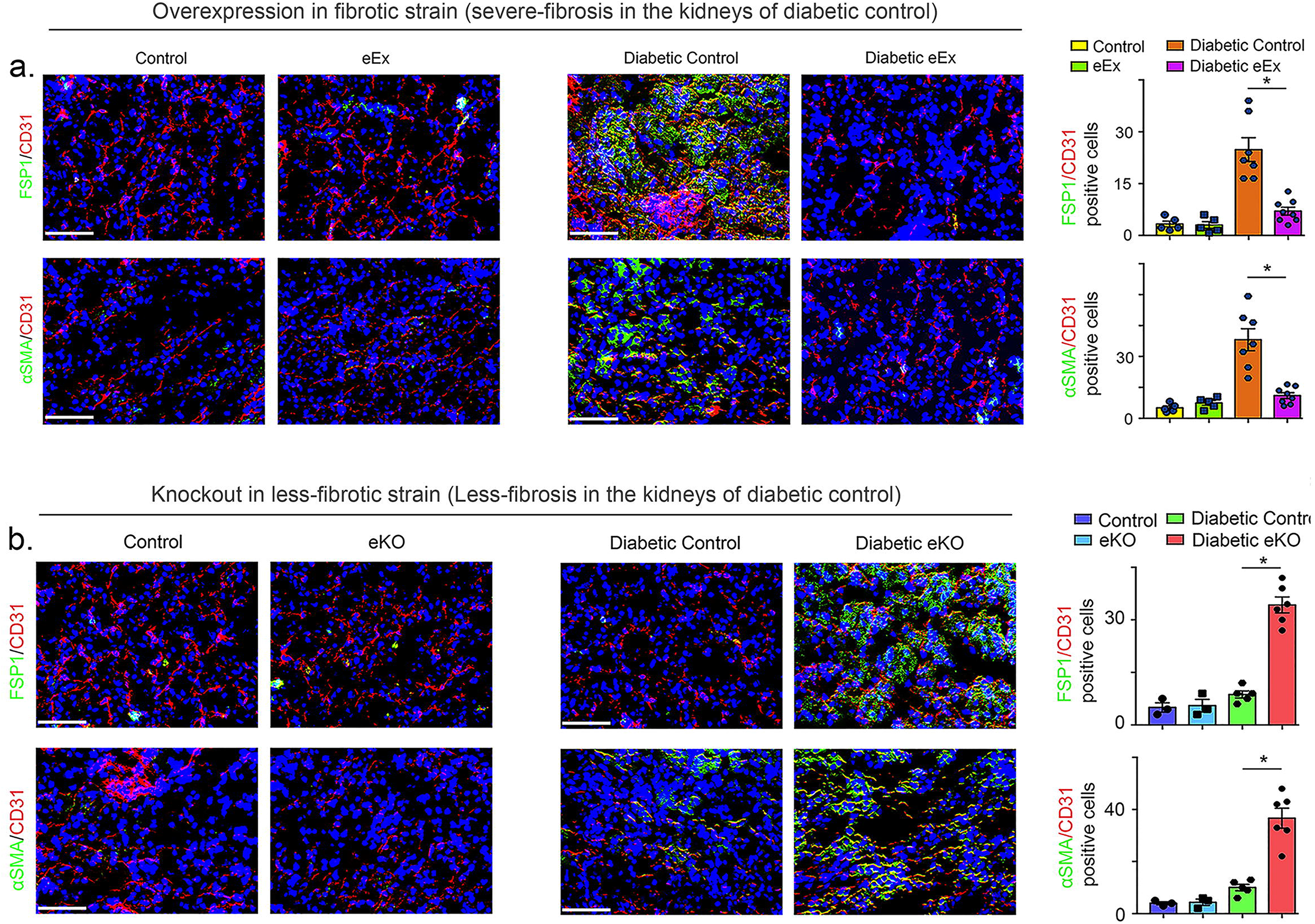
SIRT3 regulates the endothelial-to-mesenchymal transition in the kidney. **(a)** Immuno-fluorescence analysis was performed in the kidneys of non-diabetic and diabetic, control littermates and eEx mice by fluorescence microscopy. FSP-1 and α-SMA protein levels were analyzed in CD31 positive cells. Merged and representative pictures are shown. N=5 non-diabetic group, N=7 diabetic control group, N=8 diabetic eEx group. **(b)** Immunofluorescence analysis was performed in the kidneys of non-diabetic and diabetic control and eKO mice by fluorescence microscopy. FSP-1 and α-SMA protein levels were analyzed in CD31 positive cells. Merged and representative pictures are shown. N=3 non-diabetic group, N=5 diabetic control group, N=6 diabetic eKO group. Scale bar: 50 mm in each panel. 40X images are shown. Data in the graph are shown as mean ± SEM.

### SIRT3 regulates metabolic reprogramming in the endothelial cells derived myofibroblasts in the diabetic kidneys

We isolated endothelial cells from the kidneys of diabetic and non-diabetic mice **(Figure 5a)**. The endothelial cells isolated from diabetic eEx exhibited suppressed levels of αSMA, TGFβR1, smad3 phosphorylation, hexokinase 2 (HK2), pyruvate kinase M2 type (PKM2), PKM2 dimer-interconversion and pyruvate dehydrogenase kinase 4 (PDK4) when compared to diabetic controls, however, endothelial cells from kidneys of diabetic eKO displayed higher levels of αSMA, TGFβR1, smad3 phosphorylation, HK2, PKM2 and PKM2 dimer-interconversion and PDK4 when compared to diabetic littermate controls **(Figure 5b-e).** The isolated cells from diabetic eEx exhibited suppressed levels of HIF1α and higher level of HIF1α hydroxylation when compared to diabetic control littermates, however, isolated cells from kidneys of diabetic eKO displayed higher levels of HIF1α and reduced level of HIF1α hydroxylation when compared to diabetic littermate controls **(Figure S8)**. The endothelial cells isolated from diabetic eEx exhibited higher level of CPT1a and PGC1α when compared to diabetic controls; however, endothelial cells from kidneys of diabetic eKO displayed suppressed levels of CPT1a and PGC1α when compared to diabetic controls **(Figure 5f-g)**. The isolated cells from diabetic eEx exhibited suppressed hexokinase, phosphofructokinase enzyme activities, lactate level whereas and elevated intracellular ATP levels when compared to diabetic control littermates, however, isolated cells from diabetic eKO displayed higher hexokinase, phosphofructokinase enzyme activities, lactate level whereas, and reduced intracellular ATP levels when compared to diabetic control littermates **(Figure S9 and 10).** While analyzing immunofluorescent staining, we observed lower expression of glycolytic enzymes in CD31-positive cells in the kidney of diabetic eEx **(Figure 6a),** and higher protein expression levels in CD31-positive cells of diabetic eKO kidney when compared to diabetic littermate controls **(Figure 6b)**.

**Figure 5.**
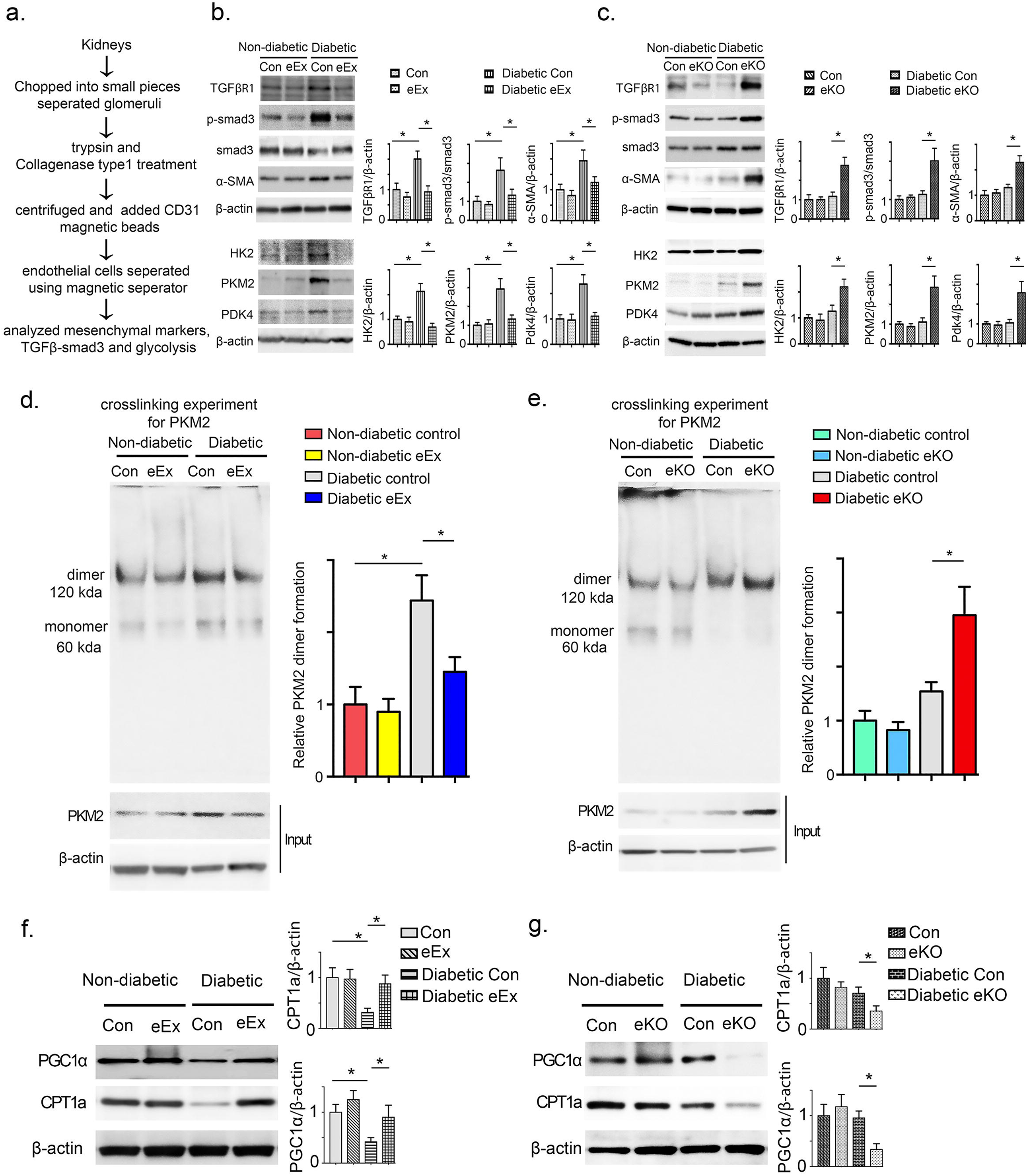
SIRT3 regulates metabolic reprogramming in the endothelial cells-derived fibroblasts in kidney. **(a)** Schematic diagram showing the isolation of endothelial cells from the non-diabetic and diabetic mice. **(b)** Western blot analysis of TGFβR1, smad3 phosphorylation, total smad3, α-SMA, HK2, PKM2 and PDK4 in the lysates of isolated endothelial cells from non-diabetic and diabetic kidneys of control littermates and eEx mice. Representative blots are shown. Densitometry calculations were normalized to β-actin. N=6 were analyzed in each group. **(c)** Western blot analysis of TGFβR1, smad3 phosphorylation, total smad3, α-SMA, HK2, PKM2 and PDK4 in the lysates of isolated endothelial cells from non-diabetic and diabetic kidneys of control and eKO mice. Representative blots are shown. Densitometry calculations were normalized to β-actin. N=5 for non-diabetic group, N=6 for diabetic group. **(d)** Glutaryldehyde chemical cross-linking experiment for PKM2, was performed in the isolated endothelial cells from the non-diabetic and diabetic kidneys of control and eEx mice. The representative blot from five blots is shown. N=5/each group. **(e)** Glutaryldehyde chemical cross-linking experiment for PKM2, was performed in the isolated endothelial cells from the non-diabetic and diabetic kidneys of control and eKO mice. The representative blot from five blots is shown. N=5/each group. **(f)** Western blot analysis of CPT1a and PGC1α in the lysates of isolated endothelial cells from the non-diabetic and diabetic kidneys of control and eEx mice. Representative blots are shown. Densitometry calculations were normalized to β-actin. N=5/group. Data in the graph are shown as mean ± SEM. **(g)** Western blot analysis of CPT1a and PGC1α in the lysates of isolated endothelial cells from the non-diabetic and diabetic kidneys of control and eKO mice. Representative blots are shown here. Densitometry calculations were normalized to β-actin. N=5 non-diabetic group, N=6 diabetic group. Data in the graph are shown as mean ± SEM.

**Figure 6.**
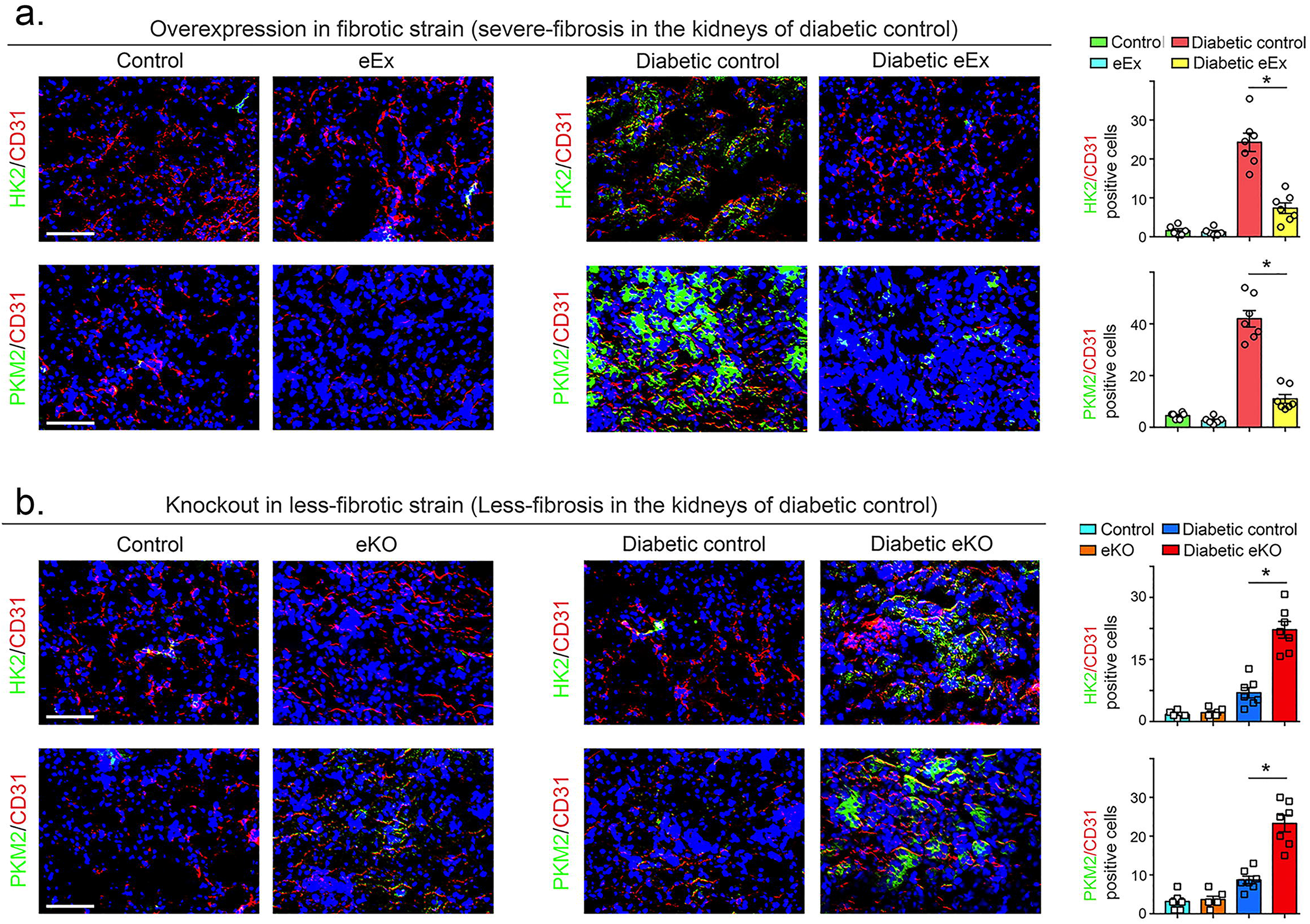
SIRT3 regulates defective glucose metabolism in the kidney endothelial cells. **(a)** Immunofluorescence analysis was performed in the kidneys of non-diabetic and diabetic control and eEx mice by fluorescence microscopy. HK2 and PKM2 protein expression was analyzed in the CD31 positive cells. Merged and representative pictures are shown. N=5 non-diabetic group, N=7 diabetic control and diabetic eEx group. **(b)** Immunofluorescence analysis was performed in the kidneys of non-diabetic and diabetic control and eKO mice by fluorescence microscopy. HK2 and PKM2 protein expression was analyzed in the CD31 positive cells. Merged and representative pictures are shown. N=5 non-diabetic group, N=7 for diabetic control and diabetic eKO group. Scale bar: 50 mm in each panel. 40X images are shown. Data in the graph are shown as mean ± SEM.

### SIRT3 deficiency disrupts metabolic homeostasis in cultured endothelial cells

To analyze the metabolic alterations specific to SIRT3, we knockdown SIRT3 in endothelial cells. While culturing the SIRT3-deplete cells in growth media, glycolysis inhibitor (dichlroacetate-DCA) and CPT1a inhibitor (etomoxir) did not alter the level of cell proliferation **(Figure 7a)**; however, culturing the SIRT3-depleted cells in diluted serum media, we observed that DCA and etomoxir caused significant reduction in the level of cell proliferation **(Figure 7a).** Moreover, we found the SIRT3-deficient cells had a higher level of glucose uptake; glycolysis inhibitors did not alter the level of glucose uptake in the SIRT3-deficient cells **(Figure 7b)**. The level of GLUT1 translocation from cytosol to cell membrane was also higher in SIRT3-depleted cells **(Figure 7c)**.

**Figure 7.**
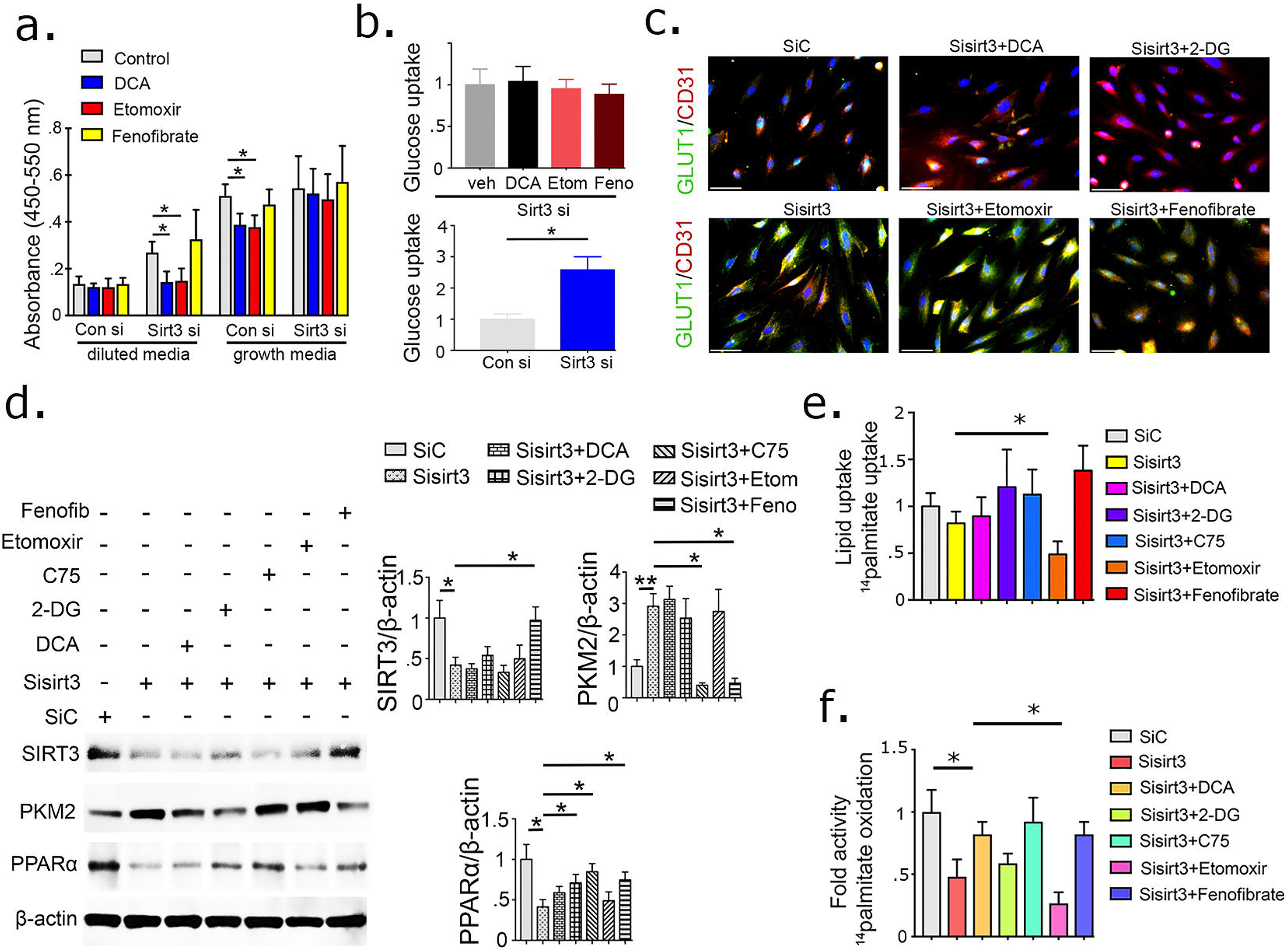
SIRT3 deficiency disrupts metabolic homeostasis in endothelial cells. **(a)** Brdu cell proliferation assay in control siRNA- and Sirt3 siRNA-transfected HUVECs in growth media and diluted media. Treatment with DCA (glycolysis inhibitor), Etomoxir (CPT1a inhibitor), and fenofibrate (PPARα agonist) in control siRNA- and SIRT3 siRNA-transfected HUVECs. Three independent sets of experiments were performed. **(b)** Glucose uptake assay in the experimental groups was analyzed by fluorimetric method. Three independent sets of experiments were performed. **(c)** GLUT1 translocation from cytoplasm to cell membrane (using CD31 as an endothelial cell marker) in the experimental groups was analyzed using immunofluorescence. GLUT1 green-FITC, CD31-red-rhodamine and DAPI-blue. Scale bar 50 μm. **(d)** Western blot analysis of SIRT3, PKM2 and PPARα in SIRT3 siRNA knockdown cells treated with glycolysis inhibitors (DCA and 2-DG), fatty acid modulators i.e. Etomoxir (CPT1a inhibitor), C75 (fatty acid synthase inhibitor) and fenofibrate (PPARα agonist). Representative blots from fours blots are shown. Densitometry analysis by Image J. The data in the each graph are normalized by β-actin. **(e)** Measurement of fatty acid uptake by radioactivity incorporation using ^14^C palmitate in 4 h serum starved HUVECs. Samples in tetraplicate were analyzed. CPM were counted and normalized to protein. **(f)** ^14^C palmitate oxidation measured by ^14^CO_2_ release. CPM were counted and normalized to the protein in the well. Samples in tetraplicate were analyzed. Data in the graph are mean ± SEM.

To probe the effect of glycolysis inhibition on TGFβ2-induced induction of mesenchymal transformation, we treated DCA with or without TGFβ2-stimulated HUVECs **(Figure S11a)**. TGFβ2 stimulation diminished the CD31 and SIRT3, while causing concomitant induction of αSMA and TGFβR1; DCA treatment restored the CD31 and SIRT3 protein levels, with significant suppression in the levels of αSMA and TGFβR1 **(Figure S11a)**. TGFβ2 did not alter the level of ^14^Cpalmitate uptake **(Figure S11b)**. TGFβ2 reduced the level of ^14^Cpalmitate oxidation; glycolysis inhibitors and FAO activations (C75) restored the level of FAO in the TGFβ2-treated cells **(Figure S11c)**.

Moreover, western blot analysis of key metabolic regulators in SIRT3 deficient cells, revealed significant suppression in PPARα; and induction in the PKM2. Glycolysis inhibition did not restore SIRT3 protein level, however, suppressed the PKM2 levels and elevated PPARα in the SIRT3 deplete cells. Fatty acid modulators (C75 and etomoxir) did not alter SIRT3 and PKM2 level whereas, fenofibrate significantly restored the level of SIRT3 and reduced the PKM2 level in the SIRT3-depleted cells **(Figure 7d).** C75 restored PPARα level in the SIRT3-depleted cells **(Figure 7d)**.

Moreover, SIRT3-depleted cells did not alter the level of lipid uptake whereas displayed significant suppression in the level of FAO **(Figure 7e-f)**. Glycolysis inhibition did not restore the level of FAO whereas, etomoxir suppressed the level of FAO in the SIRT3-deficient cells. C75 restored the suppression in FAO **(Figure 7e-f)**, suggesting that SIRT3 deficient endothelial cells reprogram the central metabolism for their survival.

### SIRT3 deficiency-linked EndMT induces mesenchymal transformation in renal tubular epithelial cells

Western blot analysis revealed that reprogrammed-SIRT3-depleted cells had induced level of αSMA, TGFβR1 and smad3 phosphorylation **(Fig. 8a and b)**. To test the contribution of reprogrammed-SIRT3 depleted cells on the mesenchymal activation in tubular epithelial cells, we transferred the conditioned media either from SIRT3 replete or SIRT3 deplete HUVECs to renal tubular epithelial cells (HK2 cells) **(Fig. 8c)**. The SIRT3 deplete cells-conditioned media (sirt3 si-CM) treatment caused significant suppression of E-cadherin levels however, caused induction of αSMA, TGFβR1 and smad3 phosphorylation protein when compared to SIRT3 replete cells-condition media (scramble si-CM) treated HK2 cells **(Fig. 8d)**. The SIRT3 deplete cells-conditioned media (sirt3 si-CM) treatment caused significant elevation in the proinflammatory IL-1β levels when compared to SIRT3 replete cells-condition media (scramble si-CM) treated HK2 cells **(Figure S12)**. To investigate whether endothelial cell SIRT3 influence EMT in the kidney, we examined E-cadherin/vimentin (EMT events) co-labeling in the diabetic kidneys. The kidneys of diabetic eEX exhibited suppressed levels of vimentin in E-cadherin positive cells when compared to diabetic controls, whereas the kidneys of diabetic eKO displayed significantly higher levels of vimentin in E-cadherin positive cells when compared to diabetic littermate controls **(Figure 8e and f)**.

**Figure 8.**
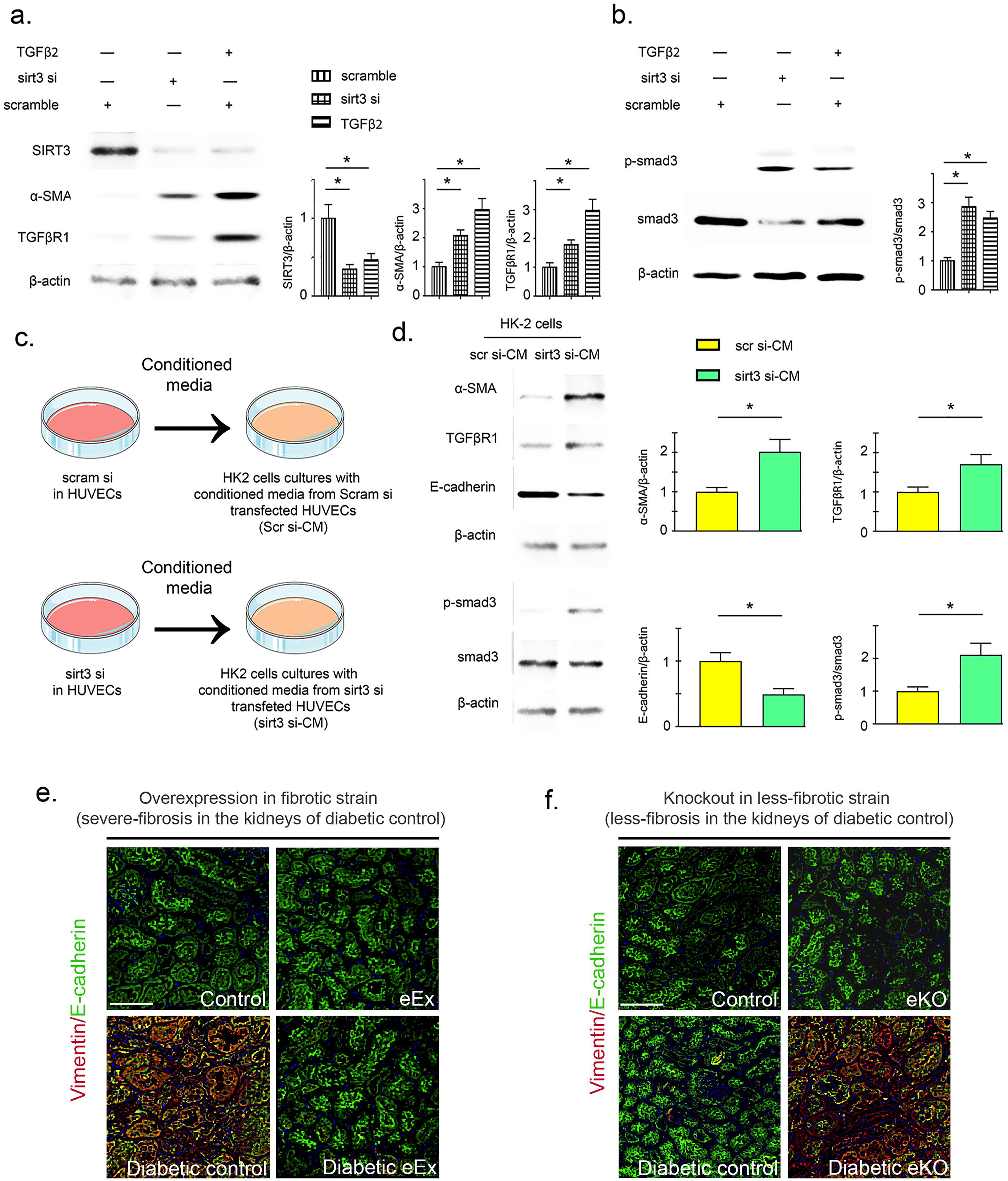
SIRT3 deficiency in endothelial cells causes mesenchymal activation in renal tubular epithelial cells. **(a)** Western blot analysis of SIRT3, α-SMA and TGFβR1 in the scramble siRNA, sirt3 siRNA-transfected and TGFβ2-stimulated HUVECs. **(b)** Smad3 phosphorylation and total smad3 in the scramble siRNA, sirt3 siRNA-transfected and TGFβ2-stimulated HUVECs. Representative blots are shown. Densitometry calculations are normalized by β-actin. Three independent experiments were performed. **(c)** Design of conditioned media experiment. Renal tubular epithelial cells (HK2 cells) were cultured in the conditioned media either from scramble siRNA- or sirt3 siRNA-transfected endothelial cells (HUVECs). **(d)** Representative western blotting images of the indicated molecules from three independent experiments are shown. Densitometric analysis of the levels relative to β-actin is shown. Data in the graph are shown as mean ± SEM. **(e)** Immunofluorescent analysis for vimentin/E-cadherin in the kidneys of nondiabetic and diabetic, control littermates and eEx mice. Representative pictures are shown. N=7/each group. Scale bar: 50 μm. **(f)** Immunofluorescent analysis for vimentin/E-cadherin in the kidneys of nondiabetic and diabetic, control littermates and eKO mice. Representative pictures are shown. N=7/each group. Scale bar: 50 μm.

## Discussion

We here, describes the crucial role of endothelial cell SIRT3 in the regulation of fibrogenic processes in the mouse model of diabetic kidney disease. We describe 1) the loss of endothelial cell SIRT3 and overexpressed level of endothelial cell SIRT3 in the renal vasculature, functions and fibrogenic processes 2) regulation of endothelial SIRT3 on central metabolic processes which affects activation of fibrogenic processes in the diabetic kidneys. Our results demonstrate that endothelial SIRT3 regulates glucose and lipid metabolism, and-associated mesenchymal trans-differentiation process by maintaining control over TGFβ-Smad3 signaling in the kidneys of diabetic mice. SIRT3 deficiency is one of the fibrotic phenotypes in diabetes that leads to PKC activation and PKM2 tetramer-to-dimer inter-conversion; ultimately, these processes alter the metabolic switch towards defective metabolism and associated mesenchymal activation in renal epithelial cells (Srivastava et al., 2018).

In addition, it is evident from our results that endothelial cell SIRT3 is critical antifibroic molecule in diabetic kidneys. Our data demonstrate that SIRT3 suppression in the kidney endothelial cells is the critical step for the metabolic reprogramming, and contributes to fibrogenic events. To test the contribution of SIRT3 in endothelial cell homeostasis, we bred the endothelial SIRT3 overexpression mouse (knock-in) and endothelial SIRT3 knock out mouse. It is clear from the results that over-expression of SIRT3 in endothelial cells attenuates fibrosis by mitigating the disrupted metabolism linked-EndMT in the kidney of diabetic mice. Our results further strengthen our views when we analyzed the loss of function of SIRT3 protein which clearly demonstrate that loss of SIRT3 in endothelial cells worsens fibrogenic processes, displays higher level of TGFβ-smad3 signaling and defective metabolism-associated-EndMT, suggesting that loss of SIRT3 disrupts the endothelial cell homeostasis, and accelerates the fibrogenic processes in the diabetic kidney. Taken together, these demonstrate that SIRT3 is critical molecule in endothelial cell metabolism and regulates EndMT in the kidney.

To envisage deeper, the contribution of defective metabolism in the endothelial cells, we used the small chemicals (which are well-known to modulate the glucose and fatty acid metabolism), in the cultured HUVECs. The results from the cultured cells confirm that SIRT3 deplete cells require a high rate of glycolysis and a level of FAO for their survival. Of note, PKM2 tetramer-to-dimer interconversion is well known regulator for central metabolism in endothelial cells (Cantelmo et al., 2016; De Bock et al., 2013; Kim et al., 2018; Schoors et al., 2014; Zhou et al., 2019). Our data suggest that SIRT3 in the endothelial cells regulates PKM2 tetramer-to-dimer interconversion and-linked disruption in central metabolism in diabetic kidney. Moreover, the data clearly demonstrate that SIRT3 deplete cells have higher GLUT1 translocation and higher glucose uptake; consequently, these processes result into the accumulation of glucose inside the cells which in turns activates HK2 enzyme in cytosol.

To begin to understand how SIRT3 deficiency-linked EndMT induce mesenchymal transformation in renal epithelial cells, we transferred the conditioned media from SIRT3 deplete endothelial cells to cultured renal tubular HK-2 cells. Interestingly, we observed the gain of mesenchymal markers, activation of TGFβ-smad3 signaling, and concomitant suppression of epithelial cell markers. These findings suggest that EndMT leads to the mesenchymal activation program (EMT) in renal tubular cells.

Current therapeutic regimens for patients who have symptom of diabetic kidney disease, include ACE inhibitors, ARBs and statins that can retard, but not prevent the progression of incidence of end-stage kidney disease in diabetes. However, side effects and intolerance to these agents often exceed their overall efficacy. The results from our data it is clear that ameliorating the level of SIRT3 in the endothelial cell can be a future strategy in combating diabetes-associated kidney fibrosis.

**Figure 9** represents the graphical representation of functional importance of SIRT3 protein in the endothelial cells homeostasis. SIRT3 deplete cells, have higher level of TGFβ-smad3 signaling, PKM2-dimer-interconversion and suppressed level of FAO and higher level of glycolysis. Cumulative effects of these metabolic changes in the SIRT3 deplete condition, results into activation of mesenchymal transformation in endothelial cells which exerts paracrine effects into neighboring tubular epithelial cells.

**Figure 9.**
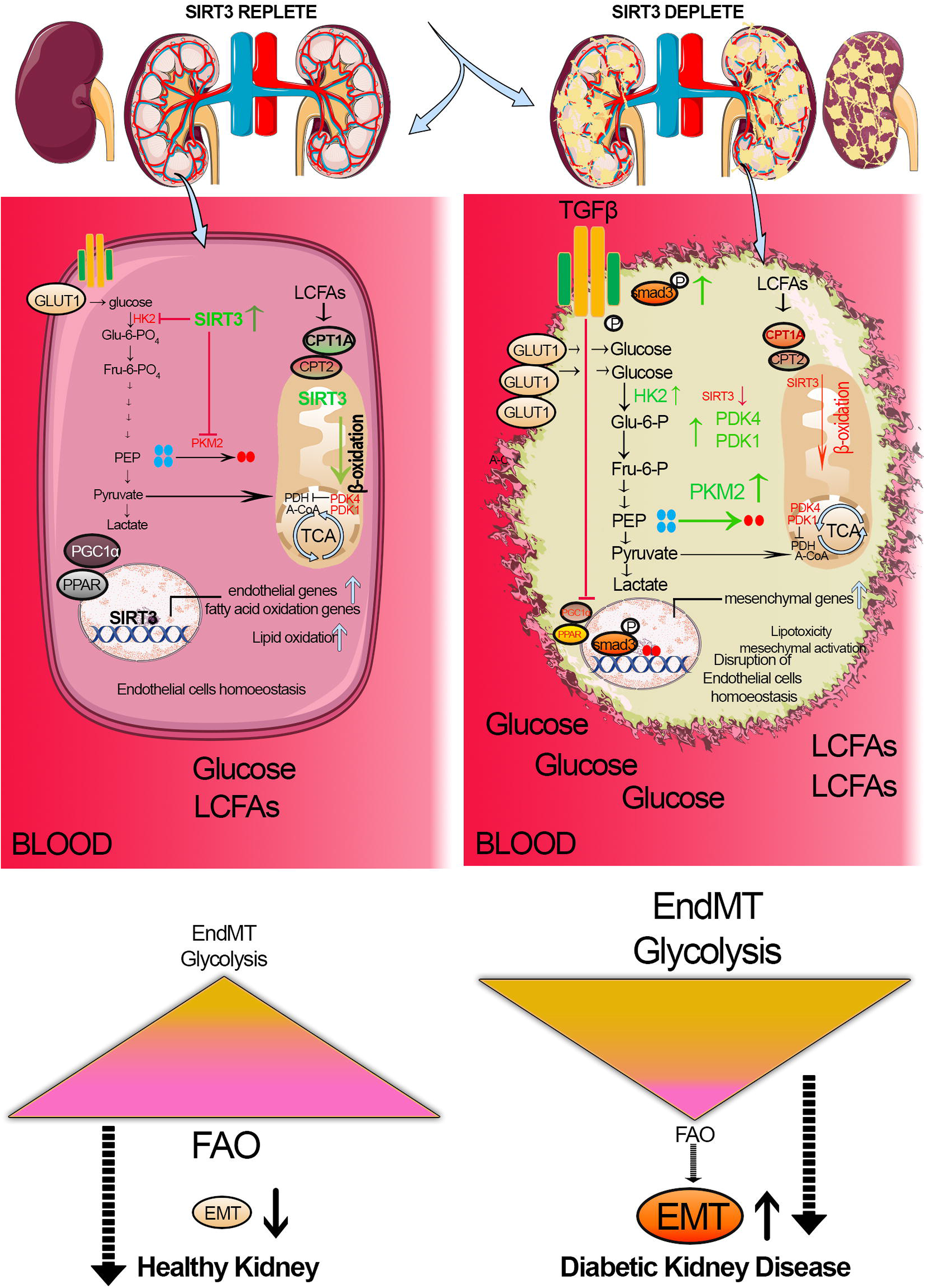
Graphical representation demonstrating the role of SIRT3 in the endothelial cell metabolism

In conclusion, our findings highlight the regulatory role of SIRT3 on EndMT in diabetic kidneys, mediated by control over TGFβ-smad3 signaling and linked-defective metabolism. SIRT3 disrupts the metabolic reprogramming in endothelial-derived-fibroblasts. This study provides new insight into the biology of SIRT3 and its regulation on endothelial cell homeostasis in the kidney.

### Limitations of study

While we observed a severe fibrosis in the diabetic kidneys of SIRT3 deficient mice (which is in the C57Bl6 mouse background); however, the study of fibrogenic phenotype in overexpression mouse (Tie1+SIRT3 tg, which is C57Bl6 mouse background) was limited since the diabetic kidneys of C57Bl6 mouse show limited fibrogenesis upon diabetes induction. Therefore, to envisage clear impact of overexpression, we bred in fibrotic mouse background (CD-1 mouse background) until 9^th^ generations. Till date, there is no suitable mouse model for diabetic kidney disease in the C57Bl6 mouse background which show severe fibrosis in kidneys. Our data highlight the regulatory role of SIRT3 in the endothelial cells however, it is still not clear how mitochondrial SIRT3 regulates glycolysis in cytosol or how cytosolic or nuclear SIRT3 regulate mitochondrial metabolism in the endothelial cells? And/or what are impact of intracellular SIRT3 translocations in the metabolic shift in endothelial cells? It will be interesting to study the role of upstream regulators of SIRT3 in different cellular compartmentations that might have a significant effect on disease phenotype in the kidney. We believe further studies will be required to understand the metabolic communications in kidney disease pathogenesis.

## STAR METHODS

### Reagent and antibodies

For rabbit polyclonal anti-pyruvate kinase (PK) isozyme M2 (4053s, RRID: AB_1904096), rabbit anti-HK-2 (2867, RRID: AB_2232946), CPT1a (D3B3, 12251) and PGC1α (3G6, 2178) antibodies were purchased from Cell Signaling Technology (Danvers, MA). A mouse monoclonal PDK4 (ab71240) were purchased from Abcam (Cambridge, UK). The goat anti-Sirt3 antibody (sc-365175, RRID: AB_10710522) was purchased from Santa Cruz Biotechnology (Dallas, TX). The mouse monoclonal anti-β-actin (AC-74) (A2228, RRID: AB_476697) antibody was obtained from Sigma (St. Louis, MO, USA). Rabbit polyclonal anti-phospho smad3 (s423 and s425) antibody was purchased from Rockland Immunochemicals (Gilbertsville, PA). Rabbit polyclonal anti-αSMA antibody (GTX100034) was purchased from GeneTex (Irvine, CA). Rabbit polyclonal anti-TGFβR1 antibody (SAB4502958) was obtained from Sigma (St Louis, MO). Fluorescence-, Alexa Fluor 647-, and rhodamine conjugated secondary antibodies were obtained from Jackson ImmunoResearch (West Grove, PA).

### Animal Experimentation

The experiments in the methods sections are carried out in accordance with Kanazawa Medical University animal protocols (protocol number 2014-89; 2013-114 and 2014-101), approved by Institutional Animal Care and Use Committee (IACUC). For the gain-of-function studies (SIRT3 over-expression), we utilized transgenic mice with over-expressed levels of SIRT3 in endothelial cells (Tie1-SIRT3 tg mice). We generated Tie1-SIRT3 tg mice in our laboratory and these mice were on the less fibrotic C57BL6 background. These mice were backcrossed for nine generations to transfer the SIRT3 tg gene into the more fibrotic CD-1 mice. We bred Tie1-SIRT3 tg+; CD-1 (eEx) and Tie1-SIRT3 tg-; CD-1 (littermate control). For the loss-of-function of SIRT3 gene, we bred the *SIRT3^flox/flox^* mice with VE-Cadherin-Cre mice (eKO) to generate mice with a deletion of the SIRT3 in their endothelial cells. *SIRT3^flox/flox^* mice and VE-Cadherin-Cre were purchased from The Jackson laboratory. The VE-Cadherin promoter used to generate the Cre-expressing mice was designed to be expressed predominantly in the endothelial cells.

The induction of diabetes in the CD-1 background mice and C57Bl6 KsJ background mice was performed according to the previously established experimental protocol (Kanasaki et al., 2014; Li et al., 2017; Nagai et al., 2014; Nitta et al., 2016; Shi et al., 2015). In brief, diabetes was induced in 10-week-old male and female eKO mice through 5 multiple consecutive doses of streptozotocin (STZ) at 50 mg/kg i.p. in 10 mmol/L citrate buffer (pH 4.5). However, a single i.p. dose of 200 mg.kg STZ was used to induce diabetes in the male and female eEx mice (eEx mice was in CD-1 mouse background, we followed previously published protocol for induction of diabetes in CD-1 mice (Kanasaki et al., 2014; Shi et al., 2015; Srivastava et al., 2016; Sugimoto et al., 2007)). Urine albumin levels were estimated using a Mouse Albumin ELISA Kit (Exocell, Philadelphia, PA).

### Morphological Evaluation

We utilized a point-counting method to evaluate the relative area of the mesangial matrix. We analyzed PAS-stained glomeruli from each mouse using a digital microscope screen grid containing 540 (27× 20) points. Masson’s trichrome-stained images were evaluated by ImageJ software, and the fibrotic areas were estimated.

### Sirius red staining

Deparaffinized sections were incubated with picrosirius red solution for 1 hour at room temperature. The slides were washed twice with acetic acid solution for 30 seconds per wash. The slides were then dehydrated in absolute alcohol three times, cleared in xylene, and mounted with a synthetic resin. Sirius red staining was analyzed using ImageJ software, and fibrotic areas were quantified.

### EndMT detection in vivo

Frozen sections (5 μm) were used for the detection of in vivo EndMT. Cells undergoing EndMT were detected by double-positive labeling for CD31 and αSMA or FSP-1 and TGFβR1. The immune-labeled sections were analyzed by fluorescence microscopy.

### Isolation of endothelial cells

Endothelial cells from the kidneys of non-diabetic and diabetic mice were isolated using the kits (Miltenyl Biotech, USA) by following the instructions from manufacturer. In general, kidneys were chopped off into small pieces and prepared for single cells suspension to carry out series of enzymatic reaction by treating with trypsin and Collagenase type I solution. The pellet is dissolved with CD31 magnetic bead and the CD31-labelled cells were separated on the magnetic separator and the cells were further purified on the column provided by manufacturer. Cell number were counted by hemocytometer and were plated on 0.1 gelatin coated Petri dishes.

### RNA isolation and qPCR

Total RNA was isolated from the isolated endothelial cells using Qiagen RNeasy Mini Kit (Qiagen, Hilden, Germany). Complementary DNA (cDNA) was generated by using the Super script (Invitrogen, Carlsbad, CA). qPCRs were performed in a 7900HT Fast real-time PCR system (Life technologies) using SYBR Green fuorescence with 10 ng of cDNA and quantifed using the delta–delta-cycle threshold (Ct) method(ΔΔCt). All experiments were performed in triplicate and 18S was utilized as an internal control. Mouse SIRT3 primers were purchased from Invitrogen.

### Chemical cross-linking

The isolated endothelial cells were lysed with RIPA lysis buffer (containing PMSF, protease inhibitor cocktail and sodium orthovanadate, which were purchased from Santa Cruz Biotechnology) for 30 minutes at 4 °C. The lysates were centrifuged at 14,000 × *g* for 15 min at 4 °C. Then, the supernatants were treated with 2.3% glutaraldehyde at a final concentration of 5% and incubated at 37 °C for 10 minutes. Tris-HCL (50 mM, pH 7.5) was used to stop the reaction. The samples were boiled with 2× sample loading buffer at 94 °C for 5 minutes and then separated by 10% SDS-PAGE.

### Western blot analysis

Protein lysates were denatured in SDS sample buffer at 100 °C for 5 min, separated on SDS-polyacrylamide gels, and blotted onto PVDF membranes (Pall Corporation, Pensacola, FL, USA) using semi-dry method. The immunoreactive bands were developed using an enhanced chemiluminescence (ECL) detection system (Pierce Biotechnology, Rockford, IL, USA) and detected using an ImageQuant LAS 400 digital biomolecular imaging system (GE Healthcare Life Sciences, Uppsala, Sweden).

### *In vitro* experiment and SIRT3 transfection

Human umbilical vein endothelial cells (HUVECs, Lonza, Basel, Switzerland) cultured in EGM medium were used in this experiment. The HUVECs cells were transfected with 100 nM of specifically designed siRNA for SIRT3 using Lipofectamine 2000 transfection reagent (Invitrogen, Carlsbad, CA, USA), according to the manufacturer’s instructions. We transfected specific SIRT3 siRNA (Invitrogen, Carlsbad, CA, USA) at a 100nM concentration in the cells. The transfected cells were treated with DCA (1mM), 2-DG (1mM), fenofibrate (1μM) and etomoxir (40 μM) for 48 hr. Glucose uptake were analyzed using the kits from Biovision Inc.

In the second set of experiments we cultured Human HK-2 cells in DMEM and Keratinocyte-SFM (1X) medium (Life Technologies Green Island NY), respectively. When the cells on the adhesion reagent reached 70% confluence, cells were cultured with conditioned media from HUVECs. The conditioned media from scramble siRNA and from SIRT3 siRNA-transfected HUVECs was collected and transferred into the HK-2 cells.

### Lipid Uptake and glucose uptake

HUVECs were incubated with medium containing 0.4 μCi [^14^C] palmitate. [^14^C] palmitate radioactivity was measured by liquid scintillation counting. Glucose uptake assay were performed using kits from Biovision, USA.

### Fatty Acid Oxidation

HUVECs were incubated with medium containing 0.75 mmol/L palmitate (conjugated to 2% fatty acid–free BSA/[^14^C] palmitate at 0.4 μCi/mL) for 2 h. 1 mL of the culture medium was transferred to a sealable tube, the cap of which housed a Whatman filter paper disc. ^14^CO2 trapped in the media was then released by acidification of media using 60% perchloric acid. Radioactivity that had become adsorbed onto the filter discs was then quantified by liquid scintillation counting.

### Statistical analysis

The data are expressed as the means ± s.e.m. The One way Anova Tukey test was performed to analyze significance, which was defined as *P* < 0.05, if not specifically mentioned. The post hoc tests were run only if F achieved P< 0.05 and there was no significant variance inhomogeneity. In each experiment, N represents the number of separate experiments (in vitro) and the number of mice (in vivo). Technical replicates were used to ensure the reliability of single values. GraphPad Prism software (Ver 5.0f) was used for the statistical analysis.

## Acknowledgements

This work was partially supported by grants from the Japan Society for the Promotion of Science for KK (23790381), DK (25282028, 25670414). This work was partially supported by a Grant for Collaborative Research awarded to DK (C2011-4, C2012-1), a Grant for Promoted Research awarded to KK (S2015-3, S2016-3, S2017-1) from Kanazawa Medical University. SPS is supported by the Japanese Government MEXT (Ministry of Education, Culture, Sports, Science, and Technology) Fellowship Program. Mitsubishi Tanabe Pharma and Ono Pharmaceutical contributed to establishing the Division of Anticipatory Molecular Food Science and Technology. KK is under the consultancy agreement with Boehringer Ingelheim. We sincerely thank Prof. Carlos Fernandez-Hernando, Vascular Biology and Therapeutic Program Yale School of Medicine for providing us necessary chemicals and materials. JG is supported by a grant from the National Institutes of Health (R01HL131952).

## Author contribution

SPS performed most of experiments, proposed the idea, contributed paper writing, generation of figures and provided intellectual input. JL managed and validated the data accuracy. MK was involved in discussion. JG provided intellectual input and experimental reagents. KK proposed experimental design, supervised the experiments, provided intellectual input, and performed final editing the manuscript. DK provided intellectual input.

## Conflicts of Interests

The authors have declared that no conflict of interest exists.

## Notes

### Competing Interest Statement

The authors have declared no competing interest.

## References

Badal, S.S., and Danesh, F.R. (2014). New insights into molecular mechanisms of diabetic kidney disease. Am J Kidney Dis 63, S63–83.

Breyer, M.D., and Susztak, K. (2016). The next generation of therapeutics for chronic kidney disease. Nat Rev Drug Discov 15, 568–588.

Cantelmo, A.R., Conradi, L.C., Brajic, A., Goveia, J., Kalucka, J., Pircher, A., Chaturvedi, P., Hol, J., Thienpont, B., Teuwen, L.A., et al. (2016). Inhibition of the Glycolytic Activator PFKFB3 in Endothelium Induces Tumor Vessel Normalization, Impairs Metastasis, and Improves Chemotherapy. Cancer Cell 30, 968–985.

Chen, P.Y., Qin, L., Barnes, C., Charisse, K., Yi, T., Zhang, X., Ali, R., Medina, P.P., Yu, J., Slack, F.J., et al. (2012). FGF regulates TGF-beta signaling and endothelial-to-mesenchymal transition via control of let-7 miRNA expression. Cell Rep 2, 1684–1696.

Chen, T., Li, J., Liu, J., Li, N., Wang, S., Liu, H., Zeng, M., Zhang, Y., and Bu, P. (2015). Activation of SIRT3 by resveratrol ameliorates cardiac fibrosis and improves cardiac function via the TGF-beta/Smad3 pathway. Am J Physiol Heart Circ Physiol 308, H424–434.

Chung, K.W., Dhillon, P., Huang, S., Sheng, X., Shrestha, R., Qiu, C., Kaufman, B.A., Park, J., Pei, L., Baur, J., et al. (2019). Mitochondrial Damage and Activation of the STING Pathway Lead to Renal Inflammation and Fibrosis. Cell Metab 30, 784–799 e785.

Cooper, M., and Warren, A.M. (2019). A promising outlook for diabetic kidney disease. Nat Rev Nephrol 15, 68–70.

Cruys, B., Wong, B.W., Kuchnio, A., Verdegem, D., Cantelmo, A.R., Conradi, L.C., Vandekeere, S., Bouche, A., Cornelissen, I., Vinckier, S., et al. (2016). Glycolytic regulation of cell rearrangement in angiogenesis. Nat Commun 7, 12240.

De Bock, K., Georgiadou, M., Schoors, S., Kuchnio, A., Wong, B.W., Cantelmo, A.R., Quaegebeur, A., Ghesquiere, B., Cauwenberghs, S., Eelen, G., et al. (2013). Role of PFKFB3-driven glycolysis in vessel sprouting. Cell 154, 651–663.

Eelen, G., de Zeeuw, P., Simons, M., and Carmeliet, P. (2015). Endothelial cell metabolism in normal and diseased vasculature. Circ Res 116, 1231–1244.

Eelen, G., de Zeeuw, P., Treps, L., Harjes, U., Wong, B.W., and Carmeliet, P. (2018). Endothelial Cell Metabolism. Physiol Rev 98, 3–58.

Grande, M.T., and Lopez-Novoa, J.M. (2009). Fibroblast activation and myofibroblast generation in obstructive nephropathy. Nature reviews. Nephrology 5, 319–328.

Grande, M.T., Sanchez-Laorden, B., Lopez-Blau, C., De Frutos, C.A., Boutet, A., Arevalo, M., Rowe, R.G., Weiss, S.J., Lopez-Novoa, J.M., and Nieto, M.A. (2015). Snail1-induced partial epithelial-to-mesenchymal transition drives renal fibrosis in mice and can be targeted to reverse established disease. Nat Med 21, 989–997.

Hershberger, K.A., Martin, A.S., and Hirschey, M.D. (2017). Role of NAD(+) and mitochondrial sirtuins in cardiac and renal diseases. Nat Rev Nephrol 13, 213–225.

Kanasaki, K., Shi, S., Kanasaki, M., He, J., Nagai, T., Nakamura, Y., Ishigaki, Y., Kitada, M., Srivastava, S.P., and Koya, D. (2014). Linagliptin-mediated DPP-4 inhibition ameliorates kidney fibrosis in streptozotocin-induced diabetic mice by inhibiting endothelial-to-mesenchymal transition in a therapeutic regimen. Diabetes 63, 2120–2131.

Kim, B., Jang, C., Dharaneeswaran, H., Li, J., Bhide, M., Yang, S., Li, K., and Arany, Z. (2018). Endothelial pyruvate kinase M2 maintains vascular integrity. J Clin Invest 128, 4543–4556.

Kizu, A., Medici, D., and Kalluri, R. (2009). Endothelial-mesenchymal transition as a novel mechanism for generating myofibroblasts during diabetic nephropathy. Am J Pathol 175, 1371–1373.

LeBleu, V.S., Taduri, G., O’Connell, J., Teng, Y., Cooke, V.G., Woda, C., Sugimoto, H., and Kalluri, R. (2013). Origin and function of myofibroblasts in kidney fibrosis. Nat Med 19, 1047–1053.

Li, J., Liu, H., Srivastava, S.P., Hu, Q., Gao, R., Li, S., Kitada, M., Wu, G., Koya, D., and Kanasaki, K. (2020). Endothelial FGFR1 (Fibroblast Growth Factor Receptor 1) Deficiency Contributes Differential Fibrogenic Effects in Kidney and Heart of Diabetic Mice. Hypertension 76, 1935–1944.

Li, J., Shi, S., Srivastava, S.P., Kitada, M., Nagai, T., Nitta, K., Kohno, M., Kanasaki, K., and Koya, D. (2017). FGFR1 is critical for the anti-endothelial mesenchymal transition effect of N-acetyl-seryl-aspartyl-lysyl-proline via induction of the MAP4K4 pathway. Cell Death Dis 8, e2965.

Liu, Y. (2011). Cellular and molecular mechanisms of renal fibrosis. Nature reviews. Nephrology 7, 684–696.

Lovisa, S., and Kalluri, R. (2018). Fatty Acid Oxidation Regulates the Activation of Endothelial-to-Mesenchymal Transition. Trends Mol Med 24, 432–434.

Medici, D. (2016). Endothelial-Mesenchymal Transition in Regenerative Medicine. Stem Cells Int 2016, 6962801.

Medici, D., and Kalluri, R. (2012). Endothelial-mesenchymal transition and its contribution to the emergence of stem cell phenotype. Semin Cancer Biol 22, 379–384.

Morigi, M., Perico, L., Rota, C., Longaretti, L., Conti, S., Rottoli, D., Novelli, R., Remuzzi, G., and Benigni, A. (2015). Sirtuin 3-dependent mitochondrial dynamic improvements protect against acute kidney injury. J Clin Invest 125, 715–726.

Nagai, T., Kanasaki, M., Srivastava, S., Nakamura, Y., Ishigaki, Y., Kitada, M., Shi, S., Kanasaki, K., and Koya, D. (2014). N-acetyl-seryl-aspartyl-lysyl-proline Inhibits Diabetes-Associated Kidney Fibrosis and Endothelial-Mesenchymal Transition. Biomed Res Int 2014.

Nitta, K., Shi, S., Nagai, T., Kanasaki, M., Kitada, M., Srivastava, S.P., Haneda, M., Kanasaki, K., and Koya, D. (2016). Oral Administration of N-Acetyl-seryl-aspartyl-lysyl-proline Ameliorates Kidney Disease in Both Type 1 and Type 2 Diabetic Mice via a Therapeutic Regimen. Biomed Res Int 2016, 9172157.

Perico, L., Morigi, M., and Benigni, A. (2016). Mitochondrial Sirtuin 3 and Renal Diseases. Nephron 134, 14–19.

Reidy, K., Kang, H.M., Hostetter, T., and Susztak, K. (2014). Molecular mechanisms of diabetic kidney disease. J Clin Invest 124, 2333–2340.

Schoors, S., Bruning, U., Missiaen, R., Queiroz, K.C., Borgers, G., Elia, I., Zecchin, A., Cantelmo, A.R., Christen, S., Goveia, J., et al. (2015). Fatty acid carbon is essential for dNTP synthesis in endothelial cells. Nature 520, 192–197.

Schoors, S., De Bock, K., Cantelmo, A.R., Georgiadou, M., Ghesquiere, B., Cauwenberghs, S., Kuchnio, A., Wong, B.W., Quaegebeur, A., Goveia, J., et al. (2014). Partial and transient reduction of glycolysis by PFKFB3 blockade reduces pathological angiogenesis. Cell Metab 19, 37–48.

Shi, S., Srivastava, S.P., Kanasaki, M., He, J., Kitada, M., Nagai, T., Nitta, K., Takagi, S., Kanasaki, K., and Koya, D. (2015). Interactions of DPP-4 and integrin beta1 influences endothelial-to-mesenchymal transition. Kidney Int 88, 479–489.

Sosulski, M.L., Gongora, R., Feghali-Bostwick, C., Lasky, J.A., and Sanchez, C.G. (2017). Sirtuin 3 Deregulation Promotes Pulmonary Fibrosis. J Gerontol A Biol Sci Med Sci 72, 595–602.

Srivastava, S.P., Goodwin, J.E., Kanasaki, K., and Koya, D. (2020a). Inhibition of Angiotensin-Converting Enzyme Ameliorates Renal Fibrosis by Mitigating DPP-4 Level and Restoring Antifibrotic MicroRNAs. Genes (Basel) 11.

Srivastava, S.P., Goodwin, J.E., Kanasaki, K., and Koya, D. (2020b). Metabolic reprogramming by N-acetyl-seryl-aspartyl-lysyl-proline protects against diabetic kidney disease. Br J Pharmacol.

Srivastava, S.P., Hedayat, F.A., Kanasaki, K., and Goodwin, J.E. (2019). microRNA Crosstalk Influences Epithelial-to-Mesenchymal, Endothelial-to-Mesenchymal, and Macrophage-to-Mesenchymal Transitions in the Kidney. Front Pharmacol 10, 904.

Srivastava, S.P., Koya, D., and Kanasaki, K. (2013). MicroRNAs in Kidney Fibrosis and Diabetic Nephropathy: Roles on EMT and EndMT. Biomed Res Int 2013, 125469.

Srivastava, S.P., Li, J., Kitada, M., Fujita, H., Yamada, Y., Goodwin, J.E., Kanasaki, K., and Koya, D. (2018). SIRT3 deficiency leads to induction of abnormal glycolysis in diabetic kidney with fibrosis. Cell Death Dis 9, 997.

Srivastava, S.P., Shi, S., Kanasaki, M., Nagai, T., Kitada, M., He, J., Nakamura, Y., Ishigaki, Y., Kanasaki, K., and Koya, D. (2016). Effect of Antifibrotic MicroRNAs Crosstalk on the Action of N-acetyl-seryl-aspartyl-lysyl-proline in Diabetes-related Kidney Fibrosis. Sci Rep 6, 29884.

Srivastava, S.P., Shi, S., Koya, D., and Kanasaki, K. (2014). Lipid mediators in diabetic nephropathy. Fibrogenesis Tissue Repair 7, 12.

Sugimoto, H., Grahovac, G., Zeisberg, M., and Kalluri, R. (2007). Renal fibrosis and glomerulosclerosis in a new mouse model of diabetic nephropathy and its regression by bone morphogenic protein-7 and advanced glycation end product inhibitors. Diabetes 56, 1825–1833.

Sundaresan, N.R., Bindu, S., Pillai, V.B., Samant, S., Pan, Y., Huang, J.Y., Gupta, M., Nagalingam, R.S., Wolfgeher, D., Verdin, E., et al. (2015). SIRT3 Blocks Aging-Associated Tissue Fibrosis in Mice by Deacetylating and Activating Glycogen Synthase Kinase 3beta. Mol Cell Biol 36, 678–692.

Theodorou, K., and Boon, R.A. (2018). Endothelial Cell Metabolism in Atherosclerosis. Front Cell Dev Biol 6, 82.

Wang, Y.Y., Jiang, H., Pan, J., Huang, X.R., Wang, Y.C., Huang, H.F., To, K.F., Nikolic-Paterson, D.J., Lan, H.Y., and Chen, J.H. (2017). Macrophage-to-Myofibroblast Transition Contributes to Interstitial Fibrosis in Chronic Renal Allograft Injury. J Am Soc Nephrol 28, 2053–2067.

Xiong, J., Kawagishi, H., Yan, Y., Liu, J., Wells, Q.S., Edmunds, L.R., Fergusson, M.M., Yu, Z.X., Rovira, II, Brittain, E.L., et al. (2018). A Metabolic Basis for Endothelial-to-Mesenchymal Transition. Mol Cell 69, 689–698 e687.

Yin, F., and Cadenas, E. (2015). Mitochondria: the cellular hub of the dynamic coordinated network. Antioxid Redox Signal 22, 961–964.

Zeisberg, E.M., Potenta, S.E., Sugimoto, H., Zeisberg, M., and Kalluri, R. (2008). Fibroblasts in kidney fibrosis emerge via endothelial-to-mesenchymal transition. J Am Soc Nephrol 19, 2282–2287.

Zeisberg, E.M., Tarnavski, O., Zeisberg, M., Dorfman, A.L., McMullen, J.R., Gustafsson, E., Chandraker, A., Yuan, X., Pu, W.T., Roberts, A.B., et al. (2007). Endothelial-to-mesenchymal transition contributes to cardiac fibrosis. Nat Med 13, 952–961.

Zeisberg, M., Hanai, J., Sugimoto, H., Mammoto, T., Charytan, D., Strutz, F., and Kalluri, R. (2003). BMP-7 counteracts TGF-beta1-induced epithelial-to-mesenchymal transition and reverses chronic renal injury. Nat Med 9, 964–968.

Zeisberg, M., and Neilson, E.G. (2010). Mechanisms of tubulointerstitial fibrosis. Journal of the American Society of Nephrology: JASN 21, 1819–1834.

Zhou, H.L., Zhang, R., Anand, P., Stomberski, C.T., Qian, Z., Hausladen, A., Wang, L., Rhee, E.P., Parikh, S.M., Karumanchi, S.A., et al. (2019). Metabolic reprogramming by the S-nitroso-CoA reductase system protects against kidney injury. Nature 565, 96–100.

