## Supplementary material for "Endothelial SIRT3 regulates myofibroblast metabolic shifts in diabetic kidneys": Online Supplemental file: Online supplemental file.pdf

**Swayam Prakash Srivastava**

**Keizo Kanasaki**

or

**Daisuke Koya**

Department of Diabetology & Endocrinology

Kanazawa Medical University

Uchinada, Ishikawa 920-0293, Japan

### Summary

Defects in endothelial cells cause deterioration in kidney function and structure. Here, we found that endothelial SIRT3 regulates metabolic reprogramming and fibrogenesis in the kidneys of diabetic mice. By analyzing, gain-of-function of the SIRT3 gene by overexpression in a fibrotic mouse strain conferred disease resistance against diabetic kidney fibrosis; while its loss-of-function in endothelial cells exacerbated the levels of diabetic kidney fibrosis. Regulation of endothelial cell SIRT3 on fibrogenic processes was due to tight control over the defective central metabolism and linked-activation of endothelial-to-mesenchymal transition (EndMT). SIRT3 deficiency in endothelial cells stimulated the TGFβ/Smad3-depandent mesenchymal transformations in renal tubular epithelial cells. These data demonstrate that SIRT3 regulates defective metabolism and EndMT-mediated activation of the fibrogenic pathways in the diabetic kidneys. Together, our findings show that endothelial cell SIRT3 is a fundamental regulator of defective metabolism regulating health and disease processes in the kidney.

### Legends to Supplementary Figures

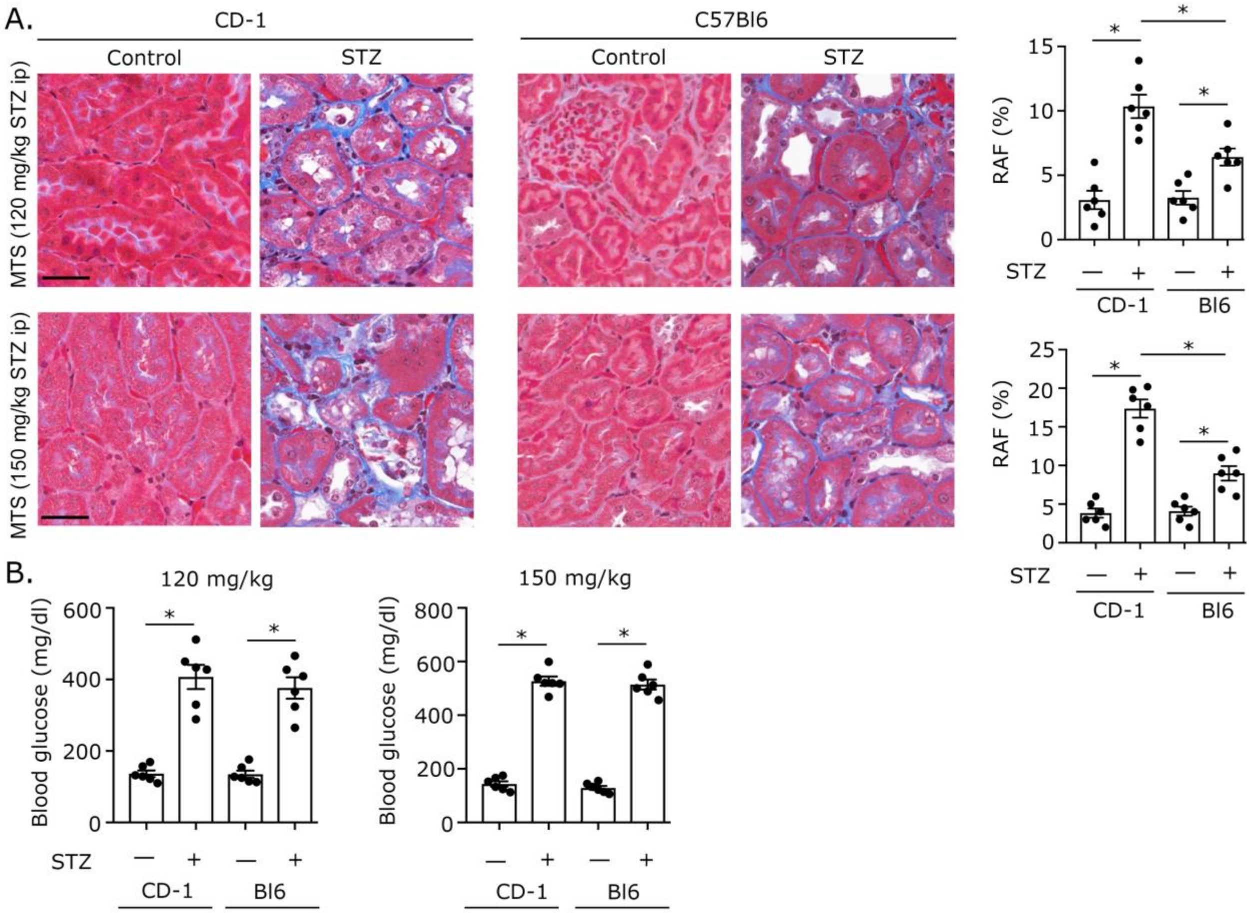


**Supplementary Figure 1 Dose dependent effect of streptozotocin on the renal fibrogenic phenotype of CD-1 and C57Bl6 mice**

**(a)** Masson trichrome staining in the kidneys of non-diabetic and diabetic CD-1 and C57Bl6 mice. Representative images are shown here. Area of fibrosis (%) was measured using the ImageJ program. N=6/each group. Data in the graph are shown as mean ± SEM. Scale bar: 50 µm.

**(b)** Blood glucose. First panel at (120 mg/kg STZ i.p. dose) and second panel at (150 mg/kg STZ i.p. dose). N=6/each group. Data in the graph are shown as mean ± SEM.

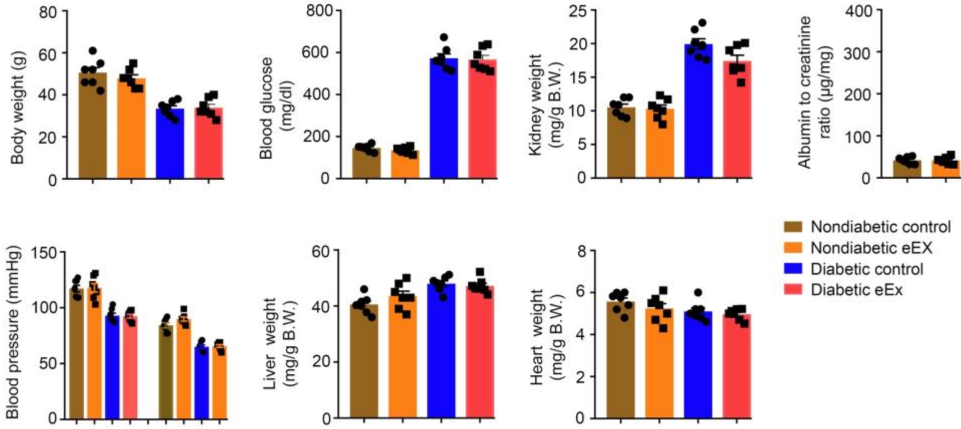


**Supplementary Figure 2 Physiological characteristics of non-diabetic and diabetic, eEx mice and littermates control**

Body weight, blood glucose, kidney weight, albumin-to-creatinine ratio (ACR), blood pressure, liver weight and heart weight were measured. N=7/each group. Data in the graph are shown as mean ± SEM.

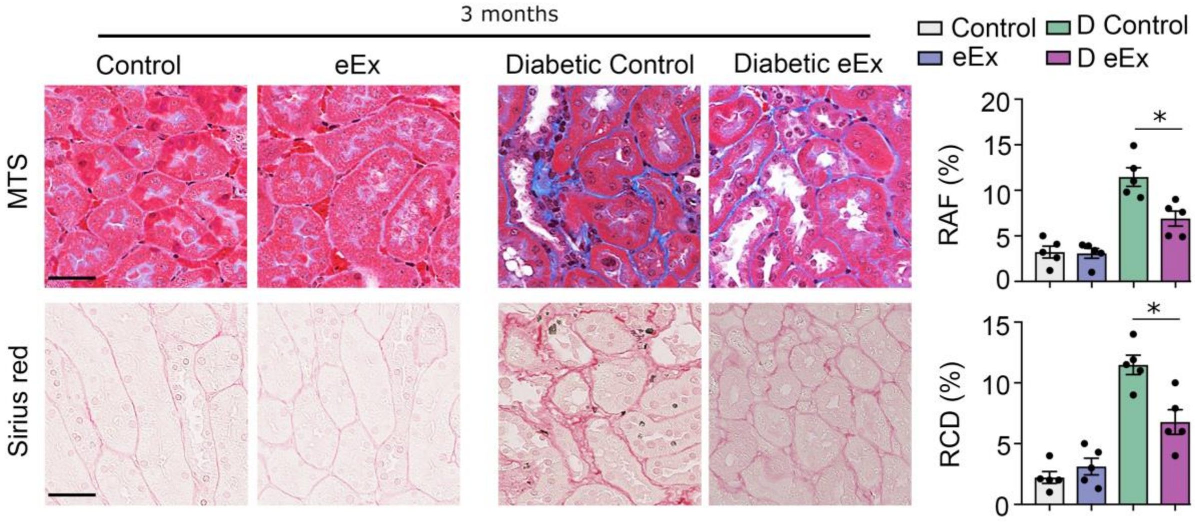


**Supplementary Figure 3 Renal fibrogenic analysis of endothelial SIRT3 overexpressed (eEx) mice and control littermates after 3 months of diabetes induction**

Masson trichrome and Sirius red staining in the kidneys of non-diabetic and diabetic eEx and control mice. Representative images are shown here. MTS-relative area of fibrosis (RAF in %) and Sirius red-relative collagen deposition (RCD in %) were measured using the ImageJ program. N=5/each group. Data in the graph are shown as mean ± SEM. Scale bar: 50 µm.

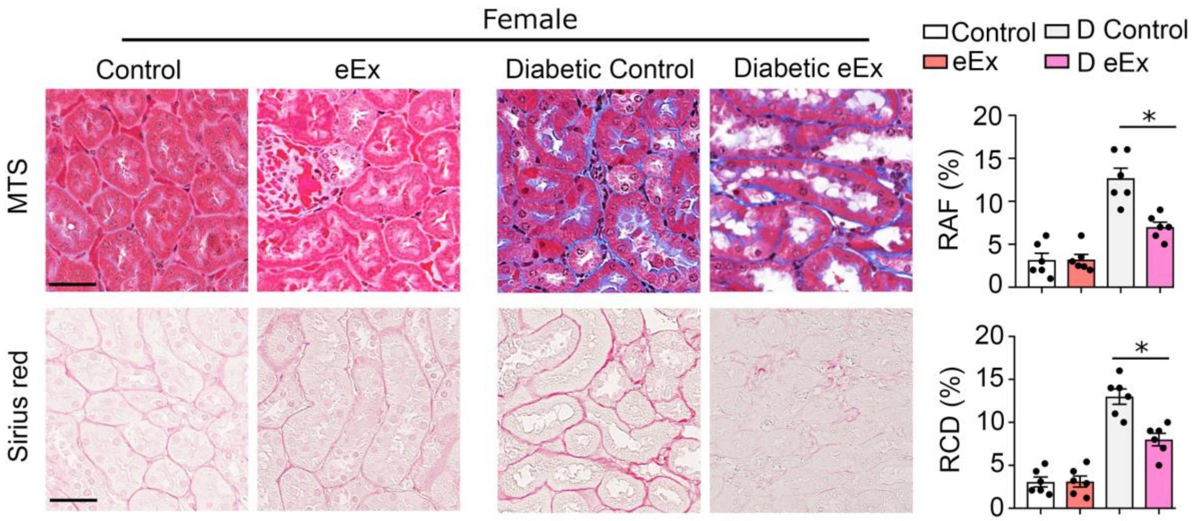


**Supplementary Figure 4 Renal fibrogenic analysis in female endothelial SIRT3 overexpressed (eEx) mice and control littermates**

Masson trichrome and Sirius red staining in the kidneys of non-diabetic and diabetic female eEx and female control mice. Representative images are shown here. MTS-relative area of fibrosis (RAF in %) and Sirius red-relative collagen deposition (RCD in %) were measured using the ImageJ program. N=6/each group. Data in the graph are shown as mean ± SEM. Scale bar: 50 µm.

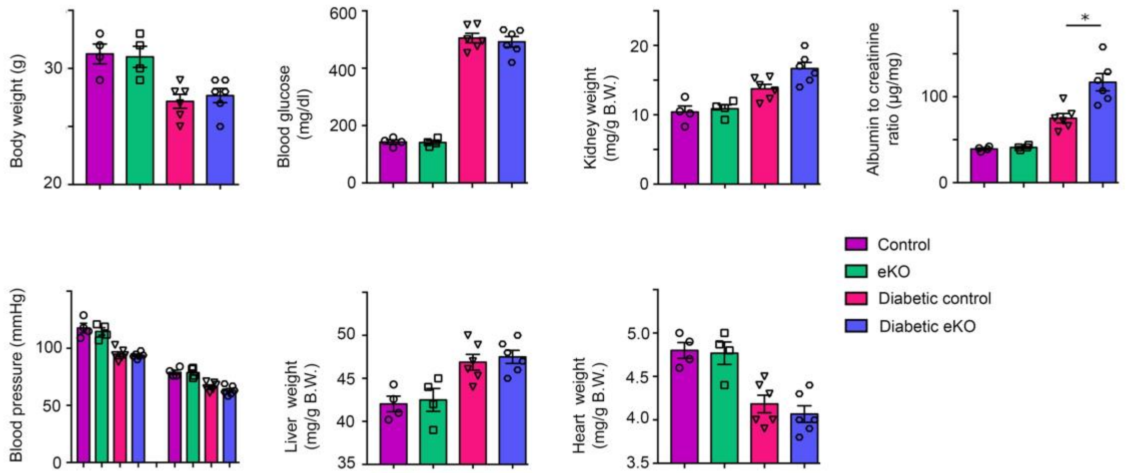


**Supplementary Figure 5 Physiological characteristics of non-diabetic and diabetic, eKO mice and control littermates**

Body weight, blood glucose, kidney weight, ACR, blood pressure, liver weight and heart weight were measured. N=4 for non-diabetic, N=6 for diabetic control and for diabetic eEx mice. Data in the graph are shown as mean ± SEM. Tukey test was used for the analysis of statistical significance.

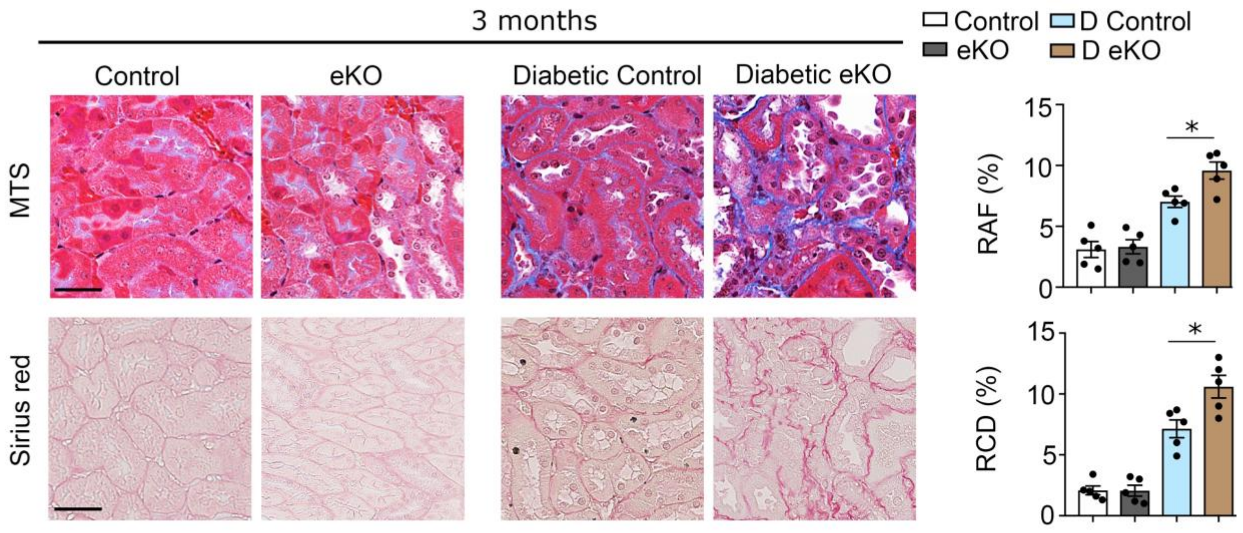


**Supplementary Figure 6 Renal fibrogenic analysis of endothelial SIRT3 knock out (eKO) mice and control littermates after 3 months of diabetes induction**

Masson trichrome and Sirius red staining in the kidneys of non-diabetic and diabetic eKO and control mice. Representative images are shown here. MTS-relative area of fibrosis (RAF in %) and Sirius red-relative collagen deposition (RCD in %) were measured using the ImageJ program. N=5/each group. Data in the graph are shown as mean ± SEM. Scale bar: 50 µm.

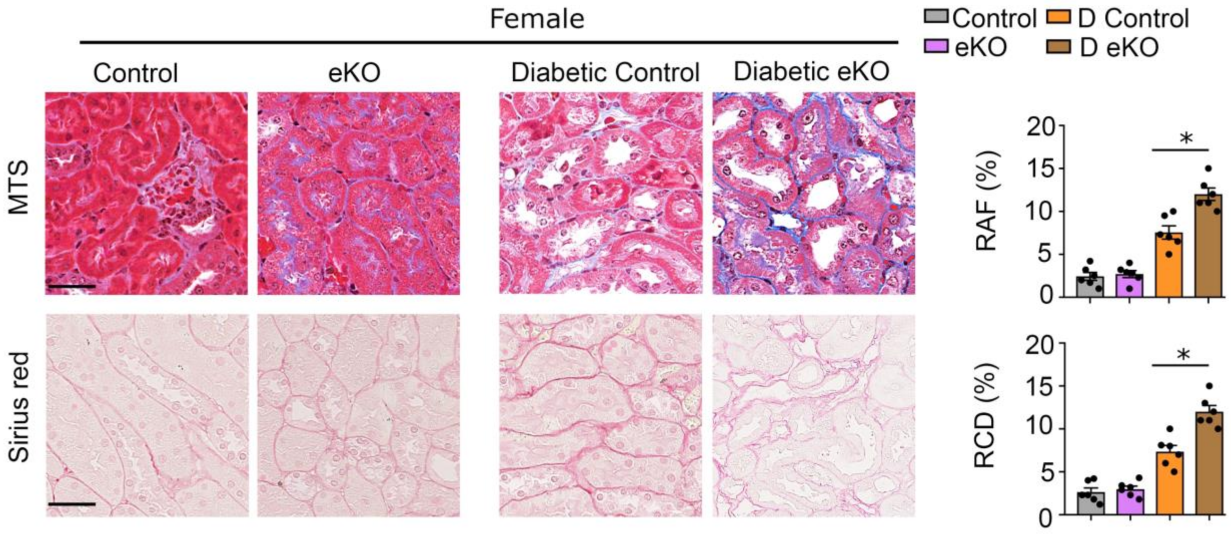


**Supplementary Figure 7 Renal fibrogenic analysis in female endothelial SIRT3 knockout (eKO) mice and control littermates**

Masson trichrome and Sirius red staining in the kidneys of non-diabetic and diabetic female eKO and female control mice. Representative images are shown here. MTS-relative area of fibrosis (RAF in %) and Sirius red-relative collagen deposition (RCD in %) were measured using the ImageJ program. N=6/each group. Data in the graph are shown as mean ± SEM. Scale bar: 50 µm.

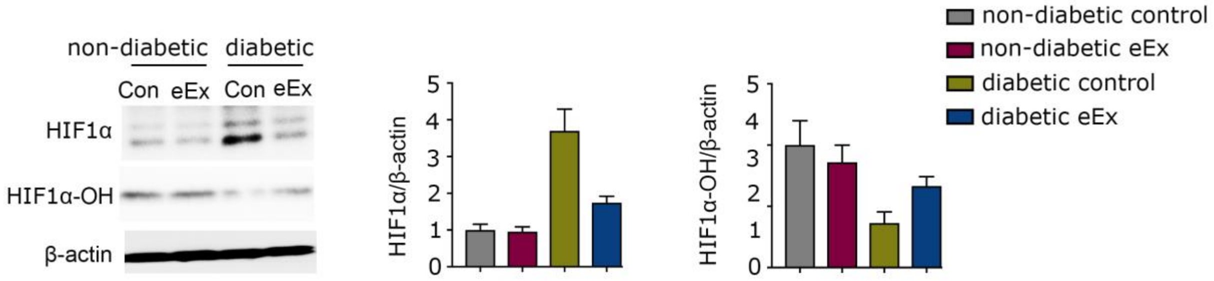


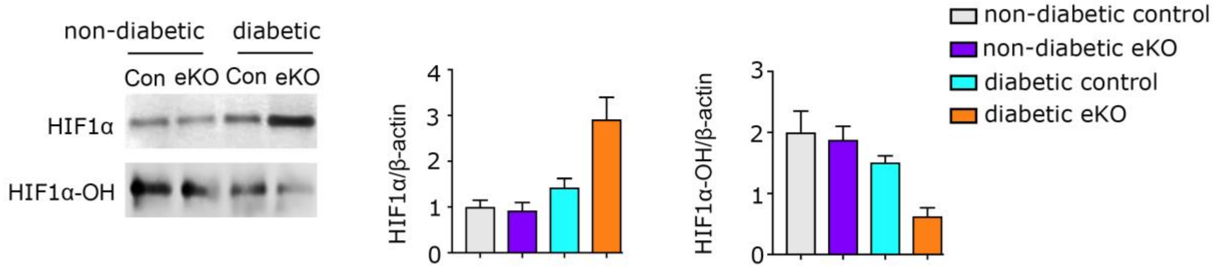


**Supplementary Figure 8 SIRT3 regulates HIF1α hydroxylation in the endothelial cells-derived fibroblasts in kidney**

First Panel- Western blot analysis of HIF1α, and HIF1-OH in the lysates of isolated endothelial cells from non-diabetic and diabetic kidneys of control and eEx mice. Representative blots are shown. Densitometry calculations were normalized to β-actin. N=6 were analyzed in each group.

Second Panel- Western blot analysis of HIF1α, and HIF1-OH in the lysates of isolated endothelial cells from non-diabetic and diabetic kidneys of control and eKO mice. Representative blots are shown. Densitometry calculations were normalized to β-actin. N=6 were analyzed in each group.

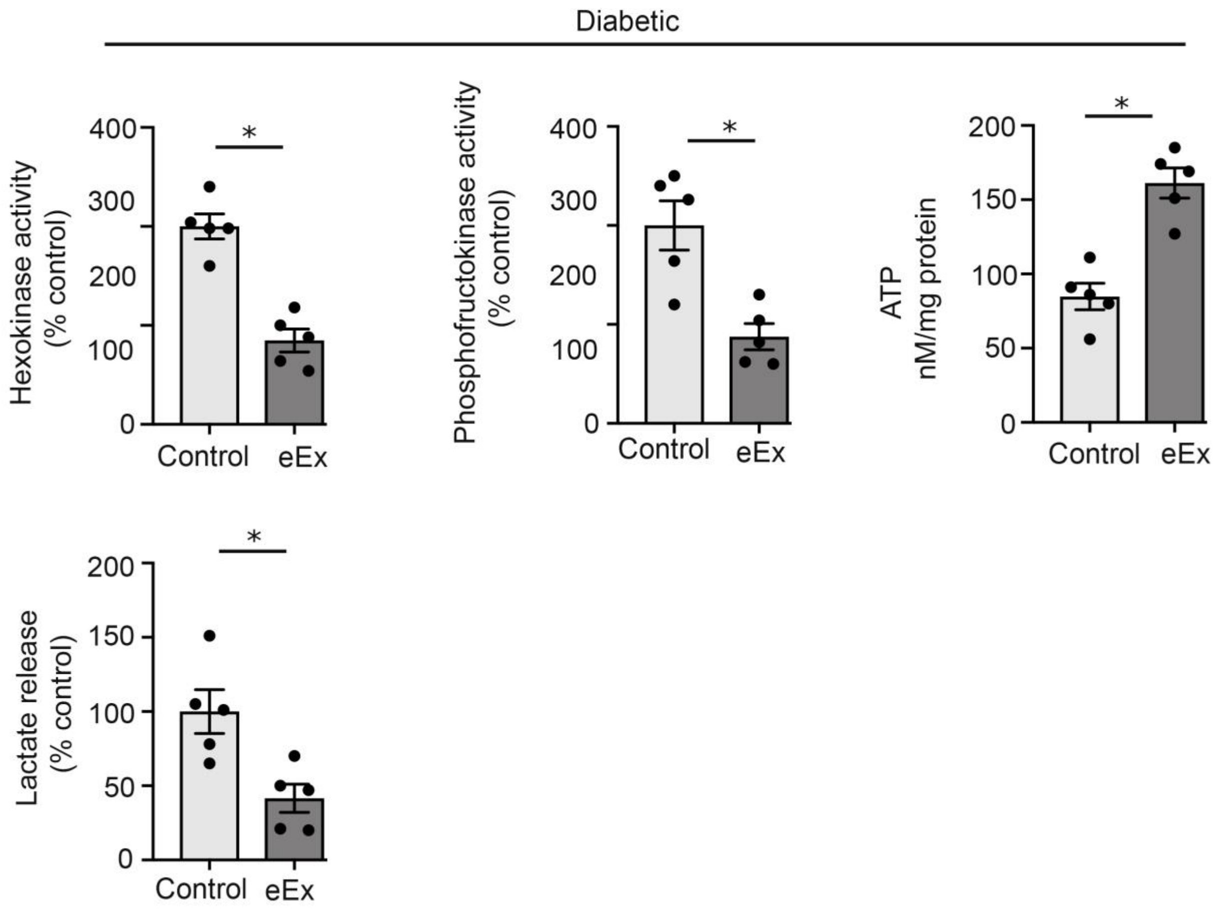


**Supplementary Figure 9 Overexpression of SIRT3 suppressed defective central metabolism in isolated endothelial cells from diabetic mice**

Hexokinase, Phosphofructokinase enzyme activities, ATP level, and lactate release in the media and PPARα transcriptional activity in the isolated endothelial cells from the diabetic control and diabetic eEx. Assays were performed using commercial kits following manufacturer's instructions. N=6/each group were analyzed. Data in the graph are shown as mean ± SEM.

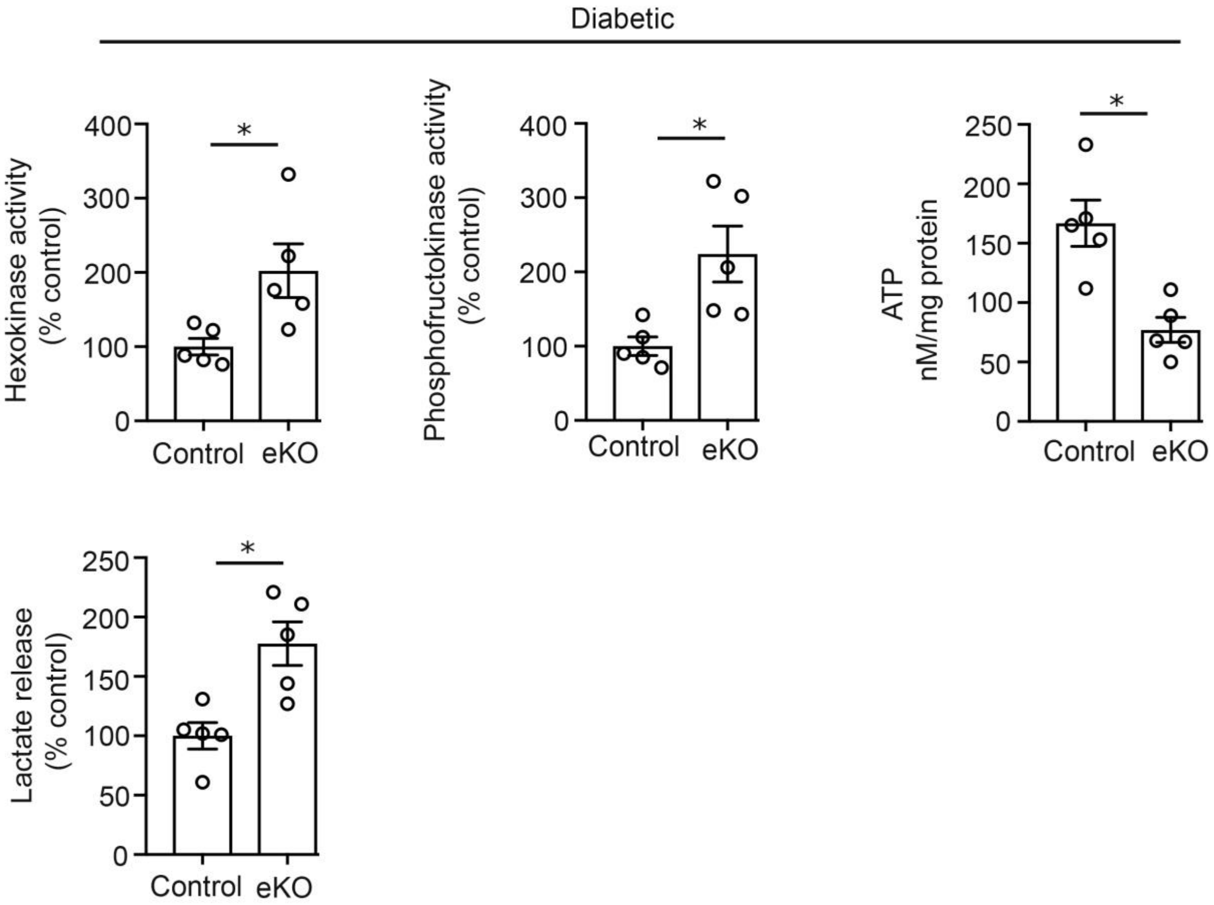


**Supplementary Figure 10 Loss of SIRT3 leads to defective central metabolism in isolated endothelial cells from diabetic mice**

Hexokinase, Phosphofructokinase enzyme activities, ATP level, and lactate release in the media and PPARα transcriptional activity in the isolated endothelial cells from the diabetic control and diabetic eKO. Assays were performed using commercial kits following manufacturer's instructions. N=5/each group were analyzed. Data in the graph are shown as mean ± SEM.

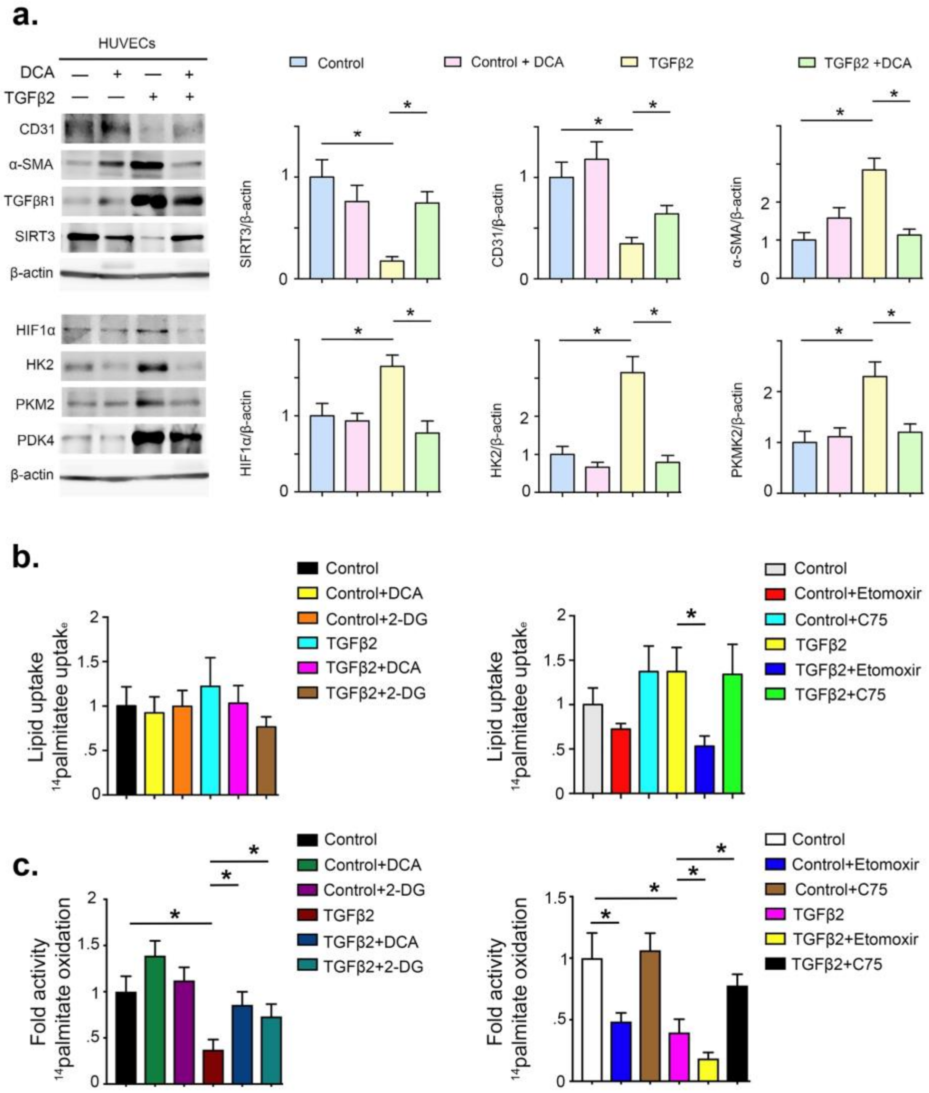


**Supplementary Figure 11 TGFβ2 causes disruption of central metabolism in endothelial cells**

**(a)** Western blot analysis of the indicated molecules in HUVEC from five independent experiments is shown. Densitometric analysis of the levels relative to β-actin is shown. Data in the graph are shown as mean ± SEM.

**(b)** Measurement of fatty acid uptake by radioactivity incorporation using [$^{14}$C]-palmitate in glycolysis inhibitors and fatty acid oxidation modulators treated with or without

TGFβ2-stimulated HUVECs. Samples in tetraplicate were analyzed. CPM were counted and normalized with protein.

(**c**) $^{14}$C palmitate oxidation by measuring $^{14}CO_2$ released. CPM were counted and normalized with protein in the well. Samples in tetraplicate were analyzed. Data in the graph are mean ± SEM. Tukey test were performed for statistical significance. Significance *-<0.05.

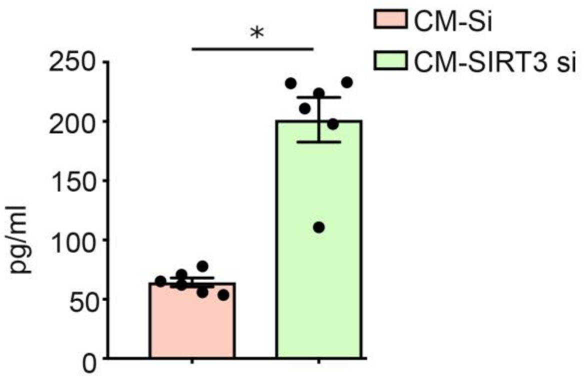


**Supplementary Figure 12 IL-1β level analysis**

Determination of IL-1β level in the indicated group. N=6/each group. Data in the graph are mean ± SEM. Tukey test were performed for statistical significance. Significance *-<0.05.
